# Genome sequences of *Aegilops* species of section Sitopsis reveal phylogenetic relationships and provide resources for wheat improvement

**DOI:** 10.1101/2021.08.09.455628

**Authors:** Raz Avni, Thomas Lux, Anna Minz-Dub, Eitan Millet, Hanan Sela, Assaf Distelfeld, Jasline Deek, Guotai Yu, Burkhard Steuernagel, Curtis Pozniak, Jennifer Ens, Heidrun Gundlach, Klaus F. X. Mayer, Axel Himmelbach, Nils Stein, Martin Mascher, Manuel Spannagl, Brande B. H. Wulff, Amir Sharon

**Author notes:** Corresponding author. (A.S), (B.B.H.W). Leibniz-Institute of Plant Genetics and Crop Plant Research (IPK) Gatersleben, Corrensstrasse 3, 06466 Seeland, Germany. Institute of Evolution, Department of Evolutionary and Environmental Biology, Faculty of Natural Sciences, University of Haifa, 199 Aba Khoushy Ave., Mount Carmel, Haifa 3498838, Israel (Hanan Sela, Assaf Distelfeld). Center for Desert Agriculture, Biological and Environmental Science and Engineering Division (BESE), King Abdullah University of Science and Technology (KAUST), Thuwal 23955-6900, Saudi Arabia.

## Abstract

*Aegilops* is a close relative of wheat (*Triticum* spp.), and *Aegilops* species in the section Sitopsis represent a rich reservoir of genetic diversity for improvement of wheat. To understand their diversity and advance their utilization, we produced whole-genome assemblies of *Ae. longissima* and *Ae. speltoides*. Whole-genome comparative analysis, along with the recently sequenced *Ae. sharonensis* genome, showed that the *Ae. longissima* and *Ae. sharonensis* genomes are highly similar and most closely related to the wheat D subgenome. By contrast, the *Ae. speltoides* genome is more closely related to the B subgenome. Haplotype block analysis supported the idea that *Ae. speltoides* is the closest ancestor of the wheat B subgenome and highlighted variable and similar genomic regions between the three *Aegilops* species and wheat. Genome-wide analysis of nucleotide-binding site leucine-rich repeat (NLR) genes revealed species-specific and lineage-specific NLR genes and variants, demonstrating the potential of *Aegilops* genomes for wheat improvement.

**Teaser:** Genome sequences of Aegilops species provides a key for efficient exploitation of this rich genetic resource in wheat improvement.

## Introduction

Wheat domestication started some 10,000 years ago in the Southern Levant with the cultivation of wild emmer wheat [WEW; *Triticum turgidum* ssp. *dicoccoides* (Körn.) Thell.; genome BBAA] (*1*). Expansion of cultivated emmer to other geographical regions, including Transcaucasia, allowed its hybridization with *Aegilops tauschii* (genome DD) and resulted in the emergence of hexaploid bread wheat (*Triticum aestivum* L. subsp. *aestivum*, genome BBAADD) (*2*). Over the next 8,500 years, bread wheat spread worldwide to occupy nearly 95% of the 215 million hectares devoted to wheat cultivation today (*3*). The success of bread wheat has been attributed at least in part to the plasticity of the hexaploid genome, which allowed wider adaptation compared with tetraploid wheat (*4*). However, further improvement of wheat is limited by its narrow genetic diversity, a result of the few hybridization events from which hexaploid wheat evolved (*5*).

To overcome the limited primary gene pool of wheat, breeders have used wide crosses with wild wheat relatives, which contain a rich reservoir of genetic diversity. Although many useful traits have been introgressed into wheat over the years (*6*), genetic constraints limit wide crosses to species that are phylogenetically close to wheat. Furthermore, some species in the secondary and tertiary gene pool of wheat (including *Ae. sharonensis* and *Ae. longissima*) cause chromosome breakage and preferential transmission of undesired gametes in wheat hybrids due to the presence of so-called gametocidal genes, which restrict interspecific hybridization (*7, 8*). Therefore, introgression requires complex cytogenetic manipulations (*9, 10*) and extensive backcrossing to recover the desired agronomic traits of the recipient wheat cultivar. These limitations can be overcome by using gene editing and genetic engineering technologies, which are not limited by plant species and can substantially expedite transfer of new traits into elite wheat cultivars (*11, 12*). However, these technologies require high-quality genomic sequences and gene annotation, which are essential for evaluation of the diversity of the respective wild species and for efficient molecular isolation of candidate genetic loci.

*Aegilops* is the closest genus to *Triticum*, and species within this genus are considered ancestors of the wheat D and B subgenomes (*13*). The progenitor of the D subgenome of bread wheat is *Ae. tauschii* (*14*), which has a homologous D genome and can be readily crossed with tetraploid and hexaploid wheat (*15*). *Aegilops* species in the section Sitopsis contain homoeologous S or S* genomes that are also closely related to wheat; however, their relationships with specific wheat subgenomes are less clear. Initially, each of the five Sitopsis members were considered potential progenitors of the wheat B subgenome (*16*), but later studies showed that *Ae. speltoides* (genome S) occupies a basal evolutionary position and is closest to the wheat B subgenome, while the other four species (genome S*) seem more closely related to the wheat D subgenome (*17,18, 19,20*). More recent analyses suggest separating *Ae. speltoides* from the other Sitopsis species, placing it phylogenetically together with *Amblyopyrum muticum* (diploid T genome) (*21,22, 23, 24*).

The *Aegilops* species in the Sitopsis section contain many useful traits, in particular for disease resistance and abiotic stress tolerance(*25, 26, 27,28*). However, because crossing of species with S and S* genomes to wheat is not straightforward, to date, only a handful of genes have been transferred from Sitopsis species to wheat. Better genomic tools and, in particular, high-quality S genome sequences will provide a deeper understanding of the genomic relationships of these species to other wheat species and enhance efforts to identify and isolate useful genes.

In this study, we generated reference-quality genome assemblies of *Ae. longissima* and *Ae. speltoides* and performed comparative analyses of these genomes together with the recently assembled *Ae. sharonensis* genome (*29*) to determine the evolutionary relationships between these Sitopsis species and wheat. Whole-genome analysis of genes encoding nucleotide-binding site leucine-rich repeat (NLR) factors, which play key roles in disease resistance, revealed species-specific and lineage-specific NLR genes and gene variants, highlighting the potential of these wild relatives as reservoirs for novel resistance genes for wheat improvement.

## Results

### Assembly details and genome alignments

We used a combination of Illumina 250-bp paired-end reads, 150-bp mate-pair reads, and 10X Genomics and Hi-C libraries for sequencing of high-molecular-weight DNA and chromosome-level assembly of *Ae. longissima* and *Ae. speltoides* genomes. Pseudomolecule assembly using the TRITEX pipeline (*30*) yielded a scaffold N50 of 3,754,329 bp for *Ae. longissima* and 3,111,390 bp for *Ae. speltoides* (table S1), and all three genomes assembled into seven chromosomes, as expected in diploid wheat. The genome of *Ae. longissima* has an assembly size of 6.70 Gb, highly similar to that of *Ae. sharonensis* (6.71 Gb) and substantially larger than the 5.13-Gb assembly of *Ae. speltoides* (Fig. 1a). These values are in agreement with nuclear DNA quantification that showed c-values of ∼7.5, ∼7.5, and ∼5.8 pg for *Ae. longissima, Ae. sharonensis*, and *Ae. speltoides*, respectively (*31*). The structural integrity of the pseudomolecules assembles of *Ae. longissima* and *Ae. speltoides* was validated by inspection of Hi-C contact matrices (fig S1). A BUSCO (*32*) analysis was conducted to assess the genome assemblies and annotation qualities. This showed a high level of genome completeness with 97.8% for *Ae. sharonensis*, 97.5% for *Ae. longissima*, and 96.4% for *Ae. speltoides* (fig S2). The predicted chromosome sizes in *Ae. longissima* and *Ae. sharonensis* were similar for all chromosomes, except for chromosome 7, which is much smaller in *Ae. longissima* due to a translocation to chromosome 4 (*33*) (Fig. 2a).

**Fig. 1.**
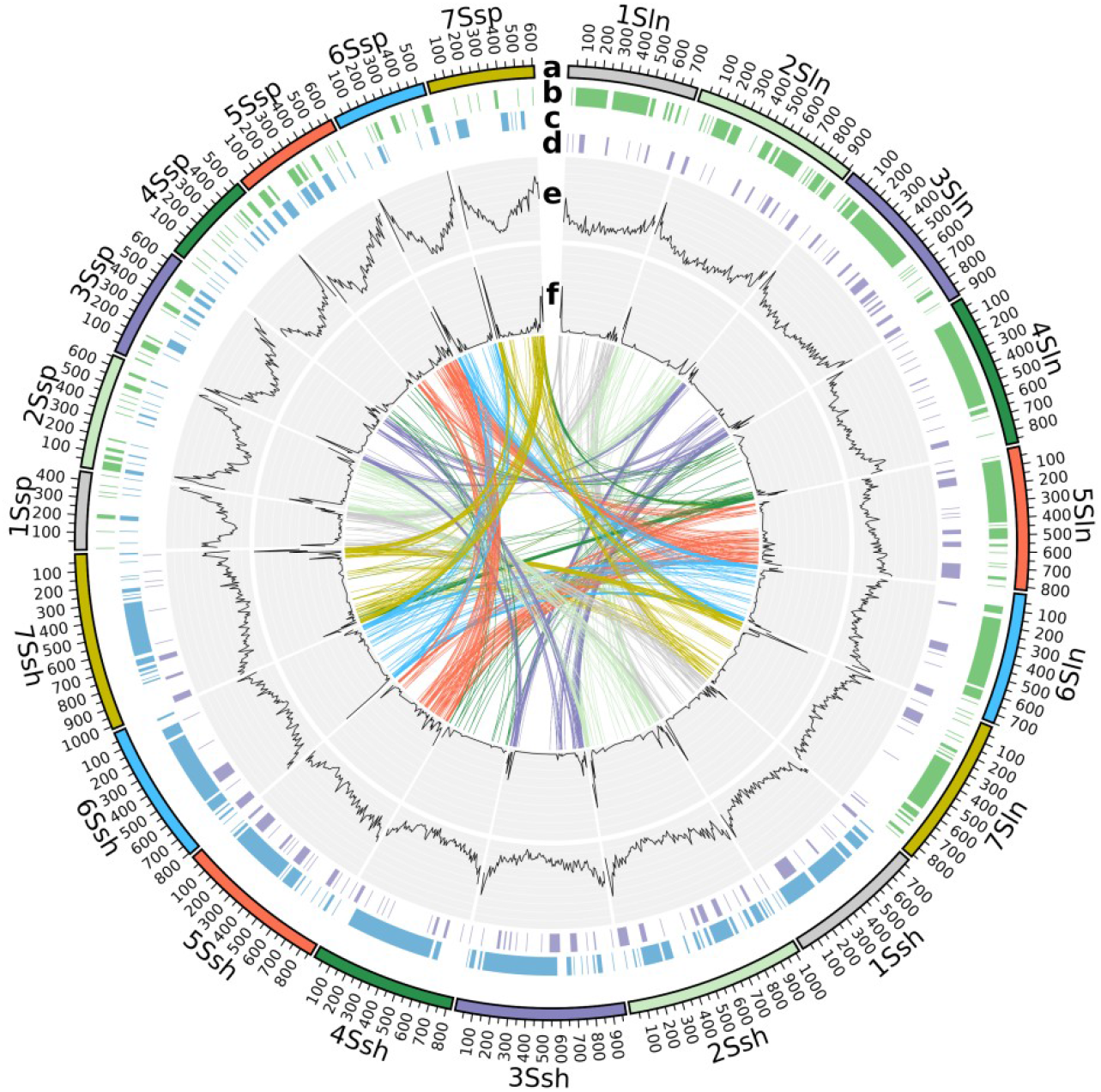
Chromosome-scale assembly and annotation of the *Ae. longissima, Ae. sharonensis*, and *Ae. speltoides* genomes. **A**, Chromosomes. **B**, Haploblocks between *Ae. sharonensis*/*Ae. longissima* and *Ae. sharonensis*/*Ae. speltoides*. **C**, Haploblocks between *Ae. longissima*/*Ae. sharonensis* and *Ae. longissima*/*Ae. speltoides*. **D**, Haploblocks between *Ae. speltoides*/*Ae. longissima* and *Ae. speltoides*/*Ae. sharonensis*. **E**, Distribution of all genes. **F**, Distribution of NLR genes. Connecting lines show links between orthologous NLR genes.

**Fig. 2.**
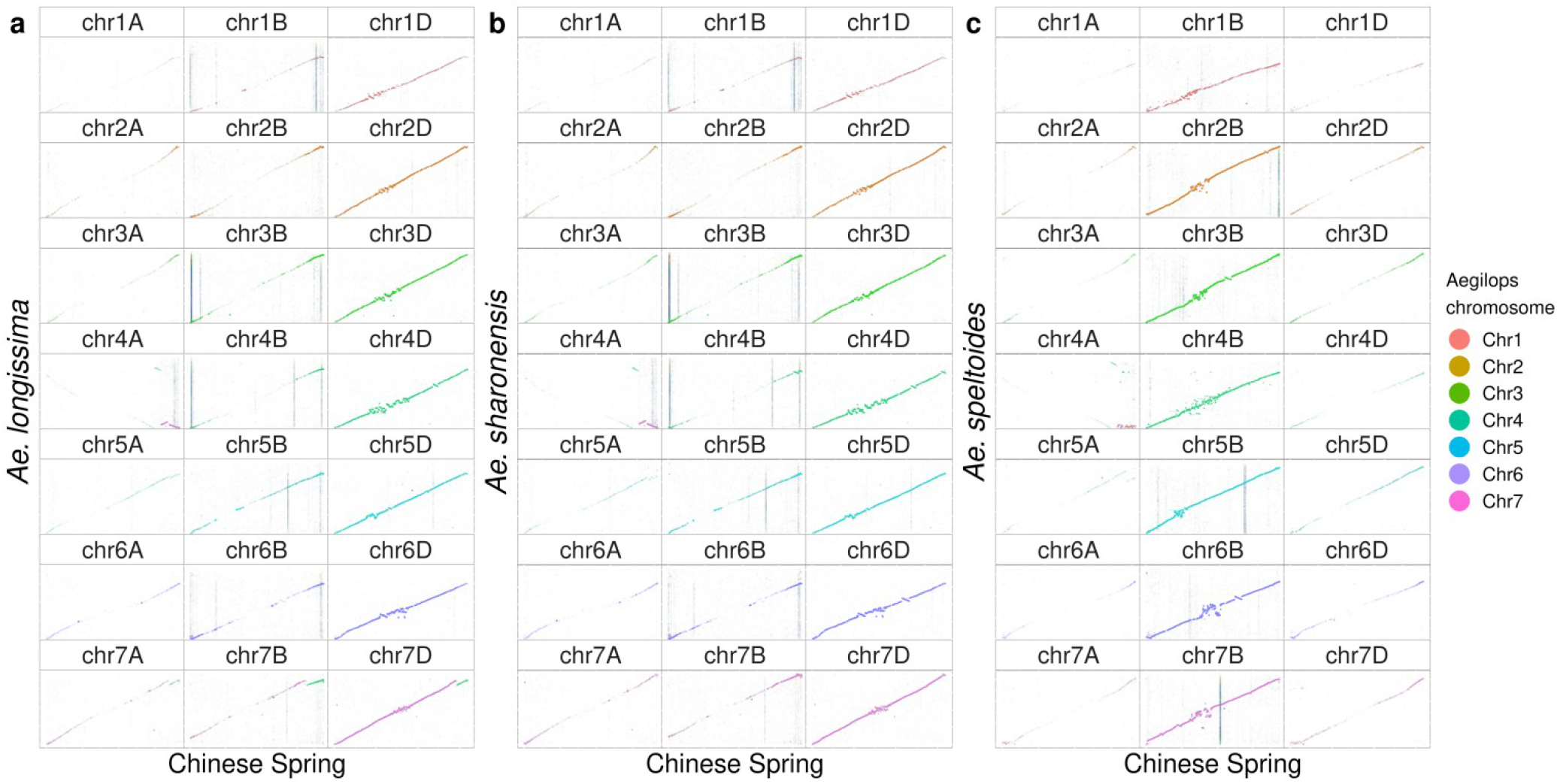
Whole-genome alignments. Alignments between *T. aestivum* cv. Chinese Spring (x-axis) and the three *Aegilops* species on the y-axis: **A**, *Ae. longissima*; **B**, *Ae. sharonensis*; **C**, *Ae. speltoides*. The best alignments are between *Ae. longissima* and *Ae. sharonensis* and the *T. aestivum* cv. Chinese Spring D subgenome and between *Ae. speltoides* and the *T. aestivum* cv. Chinese Spring B subgenome.

### Gene space annotation and ortholog analysis

We generated gene annotations for *Ae. longissima* and *Ae. speltoides* based on sequence analysis and homology to other plant species and used the same approach with the addition of RNA-seq data to generate gene annotation for the recently sequenced *Ae. sharonensis* assembly (*29*). Details of all annotations are presented in table S1. The high-confidence gene count in *Ae. speltoides* is 36,928 genes, ∼20% higher than the estimated 31,183 genes in *Ae. longissima* and 31,198 genes in *Ae. sharonensis*. The gene density in all three genomes is higher near the telomeres (Fig. 1e), similar to what has been observed in other *Triticeae* species(*14, 34, 35*). Since *Ae. speltoides* reproduces by cross-pollination, the higher number of predicted genes compared to the other two species may reflect a higher degree of heterozygosity. This notion is supported by the relatively large number of *Ae. speltoides* genes that are found on scaffolds that were not assigned to a chromosome (table S1).

We applied a whole-genome alignment approach to compare the three *Aegilops* species and the hexaploid wheat cv. Chinese Spring (CS). *Ae. longissima* and *Ae. sharonensis* both showed the best alignment to the wheat D subgenome, whereas the *Ae. speltoides* genome had the best alignment to the wheat B subgenome, and in both cases the alignment was linear (Fig. 2). To further address the phylogenetic placement of the three *Aegilops* species, we analyzed their high-confidence genes together with high-confidence genes from *Ae. tauschii* (a descendant of the donor of the D subgenome), and the subgenomes of CS (A, B, and D) and WEW (A and B). Using OrthoFinder software (*36*) (table S2), we determined orthologous groups across the gene sets and computed a consensus phylogenomic tree based on all clusters. This analysis showed the evolutionary proximity of the *Ae. speltoides* genome to the wheat B subgenome (Fig. 3) and the proximity of *Ae. longissima* and *Ae. sharonensis* to *Ae. tauschii* and the wheat D subgenome. These findings further demonstrate the evolutionary relationship between *Ae. speltoides* and the B genome, establishing it as the closest known relative of the wheat B subgenome.

**Fig. 3.**
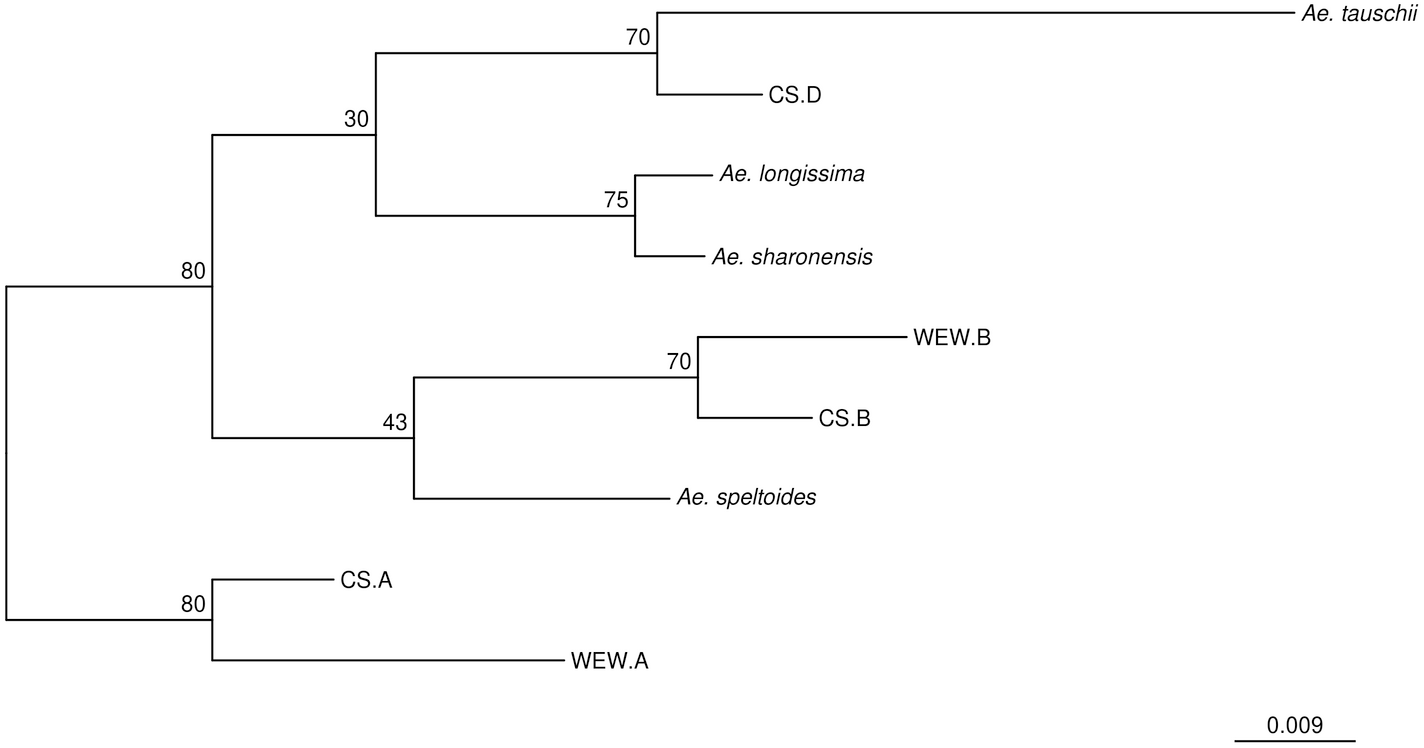
Phylogenetic tree based on OrthoFinder analysis of all high-confidence genes. Branch values correspond to OrthoFinder support values. WEW, wild emmer wheat; CS, Chinese Spring bread wheat.

We used the UpSetR package (http://gehlenborglab.org/research/projects/upsetr/) in R to analyze and visualize the number of orthogroups shared between the *Aegilops* species and the wheat subgenomes or the ones that are unique to a single species or subgenome. The “Upset plot” of the orthogroups (Fig. 4a) showed a relatively high number of shared orthogroups (292) between *Ae. longissima* and *Ae. sharonensis*. Across the B genome group (*Ae. speltoides*, and the WEW and CS B subgenomes) there were 184 orthogroups, and only 95 orthogroups were shared between the D genome group (*Ae. longissima, Ae. sharonensis, Ae. tauschii*, and the CS D subgenome). These orthologous groups contain potential candidates for group-specific genes or distinct fast-evolving genes, and their higher number in the B genome group reflects the known evolutionary distances and time scales in wheat.

**Fig. 4.**
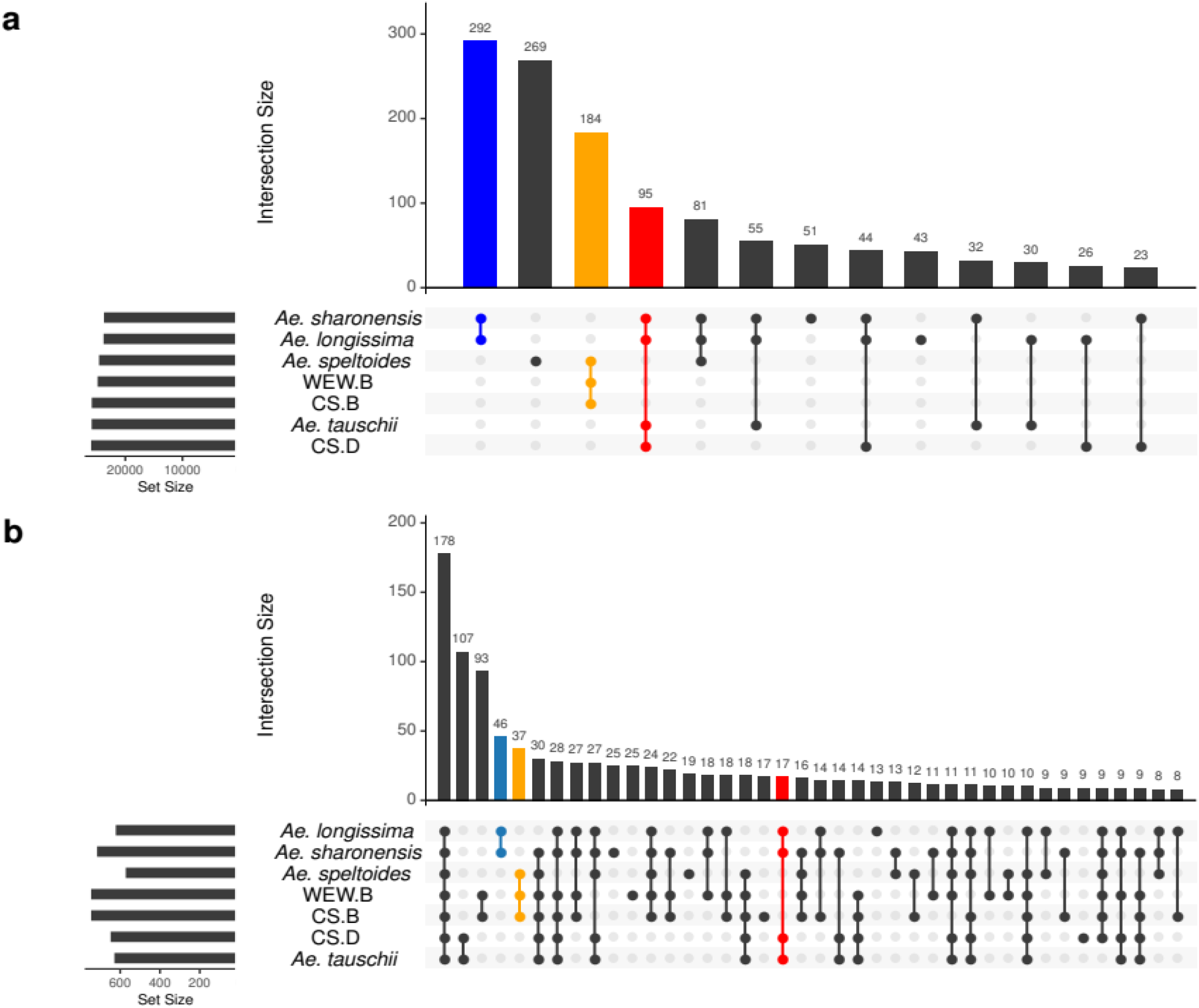
Composition of shared and unique orthogroups between different plant species and subgenomes. The species/subgenome combination is shown by the dots on the bottom panel. The bar plot shows the number of orthogroups (from the OrthoFinder analysis) for the different combinations. The histogram shows the total number of orthogroups per species/subgenome. The blue bar highlights the relatively large number of orthogroups shared by *Ae. longissima* and *Ae. sharonensis*; the orange bar shows orthogroups shared between *Ae. speltoides* and the CS and WEW B subgenomes; the red bar shows orthogroups with genes shared between *Ae. longissima, Ae. sharonensis, Ae. tauschii*, and the CS D subgenome. **A**, Orthogroups of all genes for single species or specific species combinations. **B**, Orthogroups of NLR genes for single species or specific species combinations. WEW, wild emmer wheat; CS, Chinese Spring bread wheat.

Gene alignment between *Ae. longissima* and *Ae. sharonensis* showed a 99% median sequence similarity, and the alignment of either species to the wheat subgenomes showed a median value of 97.3% for the D subgenome genes and 96.8% for the B subgenome genes. The alignment of *Ae. speltoides* high-confidence genes to the CS and WEW B genome genes had a median value of 97.3%, similar to that of *Ae. longissima* and *Ae. sharonensis* and the wheat D subgenome genes (fig S3). The complete gene annotation provided here and its relationship to wheat will be a useful resource to locate candidate genes and to target specific genes or gene families for research purposes and for breeding.

### Analysis of haplotype blocks between *Aegilops* and wheat species

We used the strategy described for wheat genomes (*37*) to construct and define haplotype blocks (haploblocks) within the *Aegilops* species as well as between wheat relatives. In our approach, we used NUCmer (*38*) to compute pairwise alignments between whole chromosomes of the respective genomes and discarded alignments smaller than 20,000 bp. We calculated the percent identity for each alignment and binned them by chromosomal position in 5-Mb bins and then combined adjacent bins sharing identical median percent identity to form a continuous haploblock. We identified haploblocks between the three *Aegilops* species (Fig. 1b-d) as well as between four crossspecies sets/combinations: (a) *Ae. sharonensis* and *Ae. longissima* (D genome relatives); (b) *Ae. longissima, Ae. sharonensis, Ae. tauschii*, and *T. aestivum* D (D genome lineage); (c) *Ae. sharonensis, Ae. longissima* (D genome relatives), and *Ae. speltoides* (B genome relative); and (d) *Ae. speltoides, T. aestivum* B, and *T. dicoccoides* B (B genome lineage) (Fig. 5). The number of identified haploblocks over all combinations varied between 53 and 93. A summary of all haploblocks and their statistics is provided as Dataset 1.

**Fig 5.**
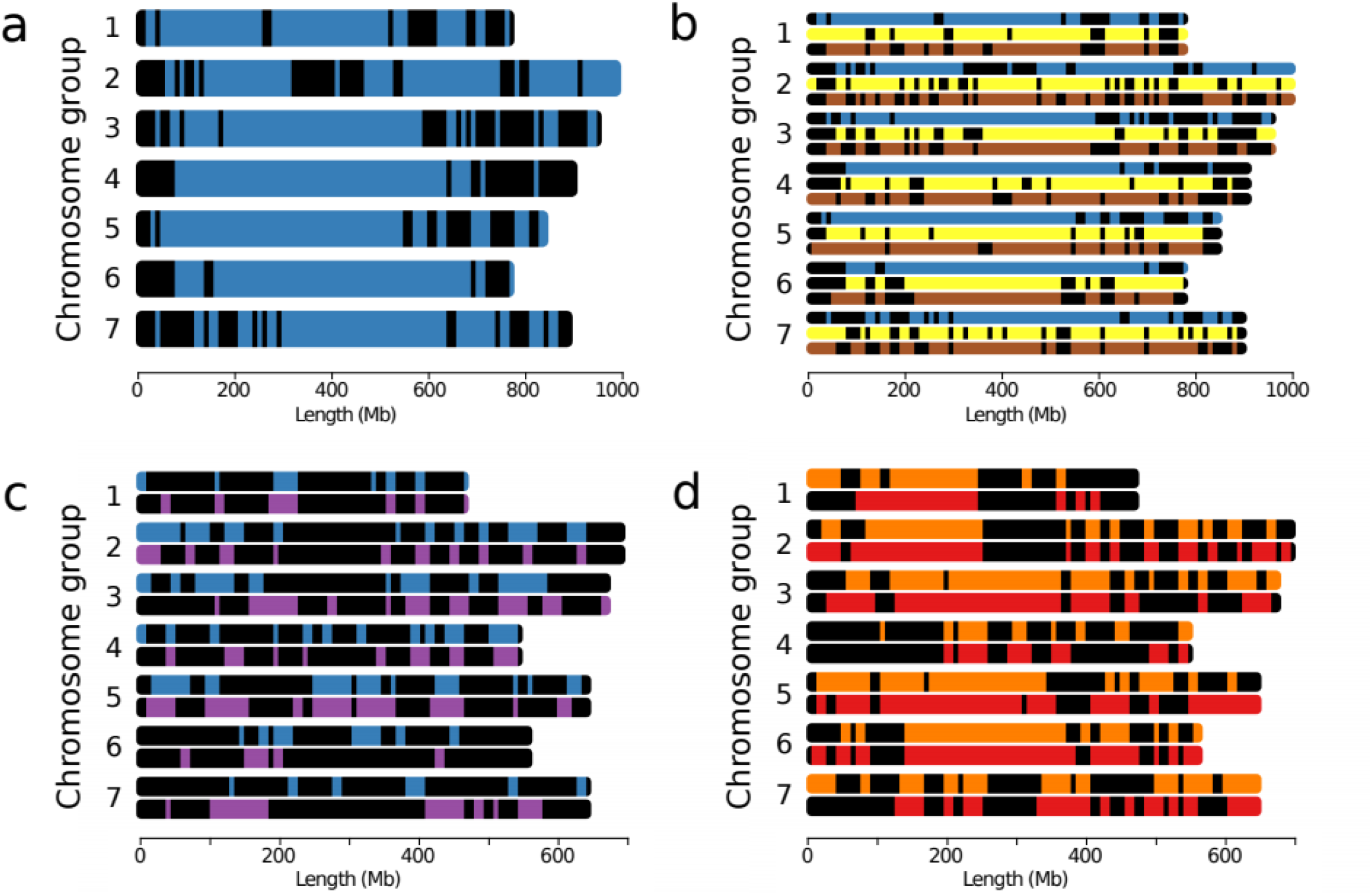
Haploblocks between *Aegilops* and wheat species. **A**, Identified haploblocks between *Ae. sharonensis* and *Ae. longissima*. Blue blocks show regions of *Ae. sharonensis* compared to *Ae. longissima* with a median identity of >99% along a 5-Mb region. **B**, Identified haploblocks between *Ae. sharonensis*, CS D, *Ae. tauschii*, and *Ae. longissima*. Blocks show regions with a median identity of >95% (>99% for *Ae. sharonensis* versus *Ae. longissima*) along a 5-Mb region. Top (blue), *Ae. sharonensis*; middle (yellow), CS D; bottom (brown), *Ae. tauschii*. **C**, Identified haploblocks between *Ae. sharonensis, Ae. longissima*, and *Ae. speltoides*. Blocks show regions with a median identity of >95% along a 5-Mb region. Top (blue), *Ae. sharonensis*; bottom (purple), *Ae. longissima*. **D**, Identified haploblocks between WEW B, CS B, and *Ae. speltoides*. Blocks show regions with a median identity of >95% along a 5-Mb region. Top (orange), CS; bottom (red), WEW. WEW, wild emmer wheat; CS, Chinese Spring bread wheat.

We identified 67 haploblocks between the highly similar *Ae. longissima* and *Ae. sharonensis* genomes (Fig. 5a), 93 haploblocks between *T. aestivum* D and *Ae. longissima*, and 90 haploblocks between *Ae. tauschii* and *Ae. longissima* (Fig. 5b). For *Ae. speltoides* and the two other *Aegilops* species, we identified 58 blocks with *Ae. sharonensis* and 53 blocks with *Ae. longissima*, with an overall good positional correlation (Fig. 5c). For the B genome relatives, we identified 63 haploblocks between *T. aestivum* B and *Ae. speltoides* and 56 between *T. dicoccoides* B and *Ae. speltoides* (Fig. 5d). In general, these blocks showed a higher positional correlation when compared to the blocks identified between the three newly sequenced *Aegilops* species. These findings indicate larger relative distance between the three *Aegilops* species under investigation with respect to conserved haploblocks compared to within the wheat B and D genome lineages.

This observation is also supported by the higher overall percentage of haploblocks covering the genome: around 20% between *Ae. speltoides, Ae. longissima*, and *Ae. sharonensis*, but more than double among the D genome relatives and among the B genome relatives. The definition of haploblocks will enable identification of diverse and similar genetic regions between the different *Aegilops* species and wheat and assist breeding efforts by allowing more targeted selection and providing convenient access to previously unused sources of genetic diversity.

### NLR repertoire in *Aegilops* spp

The Sitopsis species, in particular *Ae. sharonensis, Ae. longissima*, and *Ae. speltoides*, show pronounced variation for resistance against major diseases of wheat (*25, 27*). Since the majority of cloned disease-resistance genes encode NLRs (*39*), we decided to catalogue all NLRs present in the different genomes, analyse their genomic distribution (Fig. 1f), and study their phylogenetic relationships in *Aegilops* and wheat. To identify the NLR complement in the different genomes, we (i) searched the existing annotations of our genome assemblies for disease-resistance gene analogues and (ii) performed a *de novo* annotation using the NLR-Annotator software (*40*) (table S3). For our final list of NLRs, we compared the results of the two types of analyses and selected only those NLRs that were found by both methods.

The list of candidate NLRs that were predicted by both methods contained 742, 800, 1,030, and 2,674 candidate NLRs in the *Ae. longissima, Ae. sharonensis, Ae. speltoides*, and CS genomes, respectively (table S3). To show the diversity of the NLR repertoire in the three *Aegilops* species, we constructed a phylogenetic tree using all of the predicted NLRs along with 20 cloned NLR-type resistance genes (Fig. 6; table S4). As expected, in most cases the nearest NLR gene to a cloned gene was from CS, but in the case of *YrU1, Sr22, Lr22a, Pm2*, and *Pm3*, we also found homologs from the three *Aegilops* species (fig S4). Two large clades were completely devoid of cloned genes (Fig. 6; clusters 2 and 5). These clades might contain NLRs that are associated with resistance to pathogens or pests to which no resistance genes have yet been cloned in wheat and its wild relatives.

**Fig. 6.**
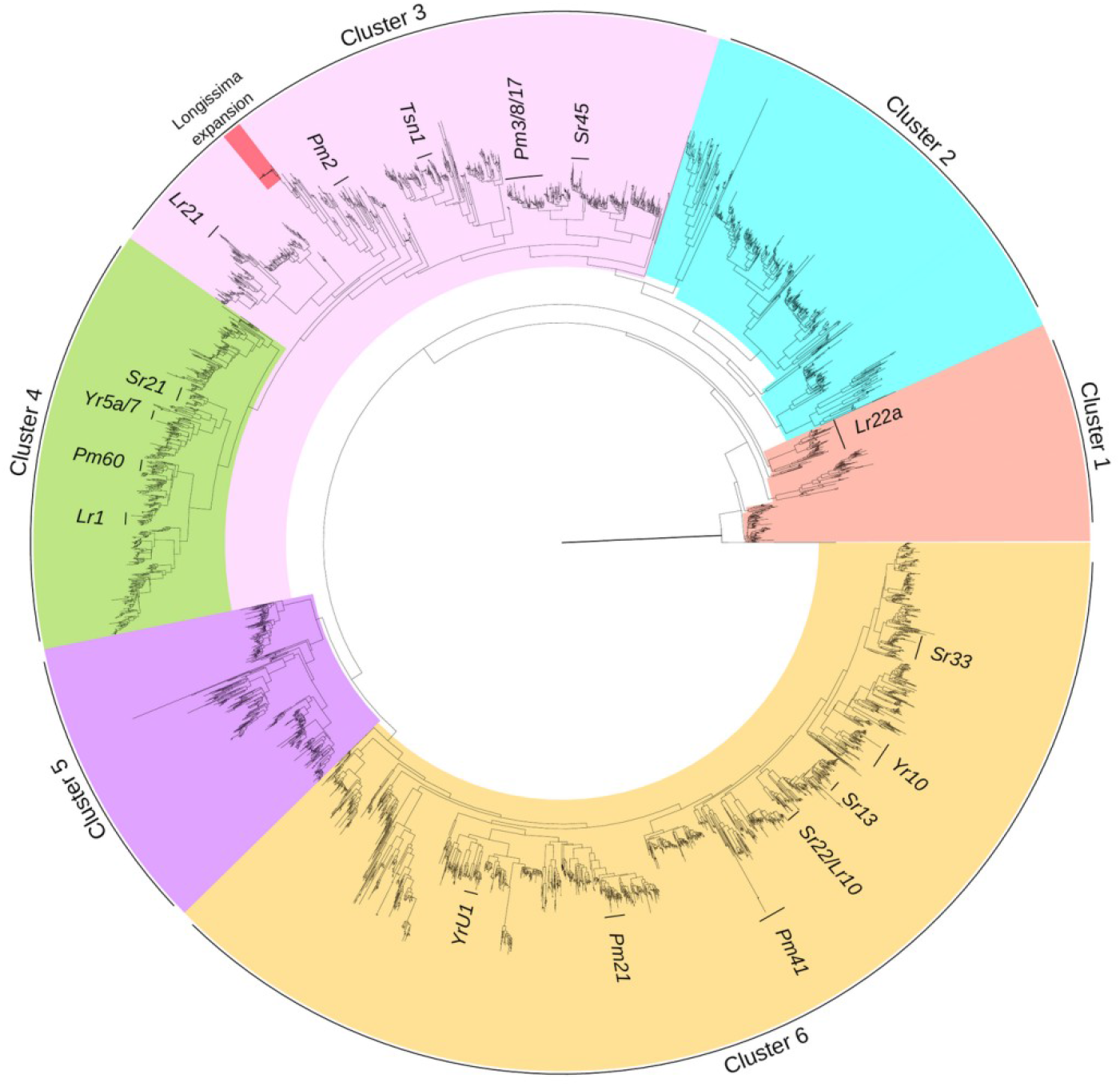
Phylogenetic tree of NLR genes in *Aegilops* and wheat. The tree illustrates NLR diversity and the genes’ similarity to cloned genes from wheat. The six clusters represent the major super-groups using an independent clustering analysis. Cluster 3 has a unique NLR expansion in *Ae. longissima* ‘chrUn’ between *Pm2* and *Lr21* containing 32 predicted genes, and this unique NLR expansion is homologous to the CS gene ‘TraesCS2B02G046000.1’ on chromosome 2B. Another expansion on *Ae. sharonensis* ‘chrUn’ that contains 11 predicted genes is located on the same branch as the *Tsn1* gene. CS, Chinese Spring bread wheat (See also Supplementary Fig 4).

We used OrthoFinder software to identify orthologous NLRs (orthoNLR) between the six genomes and found 1,312 orthoNLR groups, of which 112 were species specific. In addition, we found 42 single-copy orthoNLRs (378 predicted genes) that have one copy of each NLR present once in each of the nine diploid genomes (we split WEW into A and B subgenomes and CS into A, B, and D subgenomes). These single-copy orthoNLRs are attractive targets for assessment of their association with resistance to specific diseases and for downstream breeding applications. An overview of the distribution of the different orthoNLR groups identified in this analysis is presented in Fig. 4b. Notably, 46 NLR groups were specific to *Ae. sharonensis* and *Ae. longissima* (Fig. 4b, blue bar), highlighting a potential reservoir of species-specific resistance genes. An additional 37 groups are specific to the *Ae. speltoides*/EmmerB/WheatB B lineage (Fig. 3b, yellow bar) and 17 NLR clusters are unique to the *Ae. sharonensis*/*Ae. longissima*/*Ae. tauschii*/WheatD D lineage (Fig. 3b, red bar), both of which likely represent B and D lineage-specific NLR genes and gene variants.

A phylogenetic tree derived from the orthoNLRs (fig S5; excluding the A subgenome of CS and WEW) is congruent with the species tree obtained from orthologous group clustering of all genes (Fig. 4) and highlights the relationships of bread wheat subgenomes and the genomes of the wheat wild relatives. Based on the orthoNLR analysis, we identified all the predicted single-copy NLRs between *Ae. sharonensis, Ae. longissima*, and *Ae. speltoides*. We found 129 single-copy orthoNLRs between the three species, out of which 57 were associated with specific haploblocks, with a tendency to cluster towards distal chromosomal regions (Fig. 5).

## Discussion

Species in the genus *Aegilops* are closely related to wheat and have high genetic diversity that can potentially be used in wheat improvement. High-quality reference genome sequences are essential for efficient exploitation of these genetic resources and can also help elucidate the evolutionary and genomic relationships of these species and wheat. To this end, we sequenced and assembled the *Ae. longissima* and *Ae. speltoides* genomes and analysed them together with the recently sequenced *Ae. sharonensis* genome (*29*).

Construction of genome assemblies using the TRITEX pipeline (*30*) resulted in high-quality pseudomolecule assemblies as confirmed by BUSCO (96–98% complete genes) and whole-genome alignment to CS. Notably, the assembled genome size of *Ae. longissima* (6.70 Gb) and *Ae. sharonensis* (6.71 Gb) is substantially larger than that of *Ae. speltoides* (5.14 Gb); the genome of *Ae. speltoides* is similar in size to the published wheat subgenomes (*34, 35, 41*) and a bit larger than the *Ae. tauschii* (4.0 Gb) genome. The relative total length of the pseudomolecule assemblies of the *Ae. speltoides* genome is similar to measurements of nuclear DNA amount (*31*). Despite the relatively large genome sizes of *Ae. longissima* and *Ae. sharonensis*, their gene numbers are similar to the number of genes in all other diploid Triticeae genomes.

Species within the genus *Aegilops* have been considered the main donors of wheat diversity. *Ae. tauschii* is the direct progenitor of the wheat D subgenome, and *Aegilops* species within the section Sitopsis (S genome) were considered potential ancestors of the B genome (*16*). Later studies suggested that *Ae. speltoides* is associated with the B genome, although it is not the direct progenitor of the wheat B subgenome (*42, 43, 44*). In contrast to earlier assessments, genomic data associated the remaining Sitopsis species with the wheat D rather than the B subgenome (*17, 18,19,20*). Our analysis of the three new *Aegilops* reference genomes supports this evolutionary model and provides conclusive and quantitative evidence for the closer association of *Ae. speltoides* to the wheat B subgenome and of *Ae. longissima* and *Ae. sharonensis* to *Ae. tauschii* and the wheat D subgenome.

The best whole-genome sequence alignment of *Ae. speltoides* to the CS B subgenome (Fig. 2) and the relatively high number of shared orthogroups between *Ae. speltoides* and the CS and WEW B subgenomes (184 shared orthogroups) place *Ae. speltoides* in a “B” lineage together with the WEW and hexaploid wheat B subgenomes, while *Ae. longissima* and *Ae. sharonensis* (292 shared orthogroups) are placed in a separate “D” lineage together with *Ae. tauschii* and the wheat D subgenome (95 shared orthogroups). Accordingly, *Ae. speltoides* should be placed in a phylogenetic group outside the Sitopsis, possibly together with *Amblyopyrum muticum* (Syn. *Aegilops mutica*; diploid T genome) (*21, 22, 23,24*). The genome sequences also reveal the extremely high degree of similarity between *Ae. longissima* and *Ae. sharonensis*; the two species have an almost identical genome size and they share 292 orthogroups, compared with only 51 and 43 *Ae. longissima-* and *Ae. sharonensis*-specific groups, respectively. In fact, the only substantial genomic rearrangement between the two genomes is the unique 4S-7S translocation in *Ae. longissima*. Despite their highly similar genomes, the two species usually occupy different habitats, but occasionally they are found in mixed populations that can result in hybrids (*45*). The accessions of *Ae. longissima* and *Ae. sharonensis* used in this study were both collected in the same region, and GBS analysis showed high similarity between accessions from this area. Accessions from regions with less species overlap show more genetic differentiation (*46*). The two species demonstrate high variability in traits, such as resistance/tolerance to biotic and abiotic stresses (*47*). The high genome similarities between *Ae. longissima* and *Ae. sharonensis* along with the high phenotypic variability can be used to facilitate identification of unique traits found in these species.

The new reference genome sequences of the three *Aegilops* species are expected to advance the study and utilization of these species for wheat improvement. To make these fully accessible as resources for targeted breeding, it is necessary to unlock their genetic diversity. To this end, we constructed whole-genome haploblocks, which facilitate localization of useful variation on a genome-wide scale (*37*). Large consistent haploblocks at 99% identity were observed between the *Ae. longissima* and *Ae. sharonensis* genomes, further confirming the high similarity of these genomes, while the number of haploblocks between the three *Aegilops* species is much smaller. These findings once again demonstrate the high association between *Ae. longissima* and *Ae. sharonensis* and further support the differentiation of *Ae. speltoides* from the Sitopsis clade. Importantly, the haploblocks reveal diverse and non-diverse regions between the genomes, which points to orthologous genes with potentially beneficial variation that can be accessed by means of wide crossing or transformation.

The three *Aegilops* species contain many attractive traits, and it is expected that the new genome sequences will facilitate cloning of desired genes. For example, the *Ae. sharonensis* genome encodes resistance against a wide range of diseases that also attack wheat (*10, 27, 48,49, 50*). To better evaluate the genetic diversity and potential of disease-resistance genes in the three species, we cataloged and analyzed their NLR complements, the major class of genes encoded by plant disease-resistance genes. A high proportion of the NLR genes mapped to the telomere regions, which are also the most differentiated regions between the genomes. However, mapping the high-confidence genes between *Ae. longissima* and *Ae. sharonensis* showed a mean identity value of 98.6%, while mapping of NLR genes showed only 87% identity, suggesting that the combined diversity of disease-resistance genes present in both genomes is substantially greater than that in each of the single genomes.

Phylogenetic analysis of the NLRs from the different genomes outlined the diversity in this class of proteins and highlighted groups of genes or specific targets for further study, for example, in the two branches of the NLR phylogeny that lacked cloned resistance genes (clusters 2 and 5, Fig. 6). Alternatively, clades rich in cloned genes (such as clusters 3, 4, and 6) can be considered evolutionary hotspots. The phylogenetic tree and the orthogroup analysis both provided a cross reference to locate orthologous NLRs in CS and the three *Aegilops* species, such as the CS *Pm2* gene, which is located on a branch that contains NLRs from all of the three subgenomes as well as from the three *Aegilops* species (fig S4). Another example is the gene expansion in *Ae. longissima* that corresponds to the CS gene “TraesCS2B02G046000.1” on chromosome 2B at position 23,020,418–23,022,244 bp on the CS genome. This locus also coincides with the *MlIW39* powdery mildew resistance locus on the short arm of chromosome 2B (*51*). This cluster of NLRs is located on chrUn in *Ae. longissima*, so its exact genomic location remains to be defined. This expansion spreads over several scaffolds; therefore, it is probably not a sequencing artifact.

Recent advances in sequencing and genomic-based approaches have greatly enhanced the identification and cloning of new genes, thus expediting sourcing of candidate genes for next-generation breeding (cloned gene table within *52, 53*). The genome sequences of the three *Aegilops* species reported here reveal important details of the genetic makeup of these species and their association with durum and bread wheat. These new discoveries and the availability of high-quality reference genomes pave the way for more efficient utilization of these species, which have long been recognized as important genetic resources for wheat improvement.

## Materials and Methods

### Plant materials

*Aegilops longissima* accession AEG-6782-2 was collected from Ashdod, Israel (31.84N, 34.70E). *Ae. speltoides* ssp. *speltoides* accession AEG-9674-1 was collected from Tivon, Israel (32.70N, 35.10E). Each accession was self-pollinated for four generations to increase homozygosity. All accessions were propagated and maintained at the Lieberman Okinow gene bank at the Institute for Cereal Crops Improvement at Tel Aviv University.

Leaf samples were collected from seedlings grown at 22 ± 2°C for 2–3 weeks with a 12-h-light/12-h-dark regime in fertile soil. Before harvesting of leaves, the plants were maintained for 48 h in a dark room to lower the amounts of plant metabolites. Samples were collected directly before extraction.

### Preparation of high-molecular-weight DNA

High-molecular-weight DNA was extracted using the liquid nitrogen grinding protocol (BioNano Genomics, San Diego) and according to (*54*) with modifications as follows. All steps were performed in a fume hood on ice using ice-cold solutions. Approximately 1 g of fresh leaf tissue was placed in a Petri dish with 4 ml nuclear isolation buffer (NIB, 10 mM Tris-HCl, pH 8, 10 mM EDTA, 80 mM KCl, 0.5 M sucrose, 1 mM spermidine, 1 mM spermine, 8% PVP) and cut into 2 × 2-mm pieces using a razor blade. The volume of NIB was brought to 10 ml, and the material was homogenized using a handheld homogenizer (QIAGEN, 9001271) for 60 s. Then, 3.75 ml of β-mercaptoethanol and 2.5 ml of 10% Triton X-100 were added, and the homogenate was filtered through a 100-µm filter (VWR, 21008950), followed by washing three times with 1 ml of NIBM (NIB supplemented with 0.075% β-mercaptoethanol). The homogenate was filtered through a 40-µm filter (VWR, 21008949), the volume of the filtrate was adjusted to 45 ml by adding NIBTM (NIB supplemented with 0.075% β-mercaptoethanol and 0.2% Triton-X 100), and the mixture was centrifuged at 2000 *g* for 20 min at 4°C. The nuclear pellet was re-suspended in 1 ml of NIBM, and NIBTM was added to a final volume of 4 ml. The nuclei were layered onto cushions made of 5 ml 70% Percoll in NIBTM and centrifuged at 600 *g* for 25 min at 4°C in a swinging bucket centrifuge. The pelleted nuclei were washed once by re-suspension in 10 ml of NIBM, centrifuged at 2000 *g* for 25 min at 4°C, washed three times each with 10 ml of NIBM, and then centrifuged at 3000 *g* for 25 min at 4°C. The pelleted nuclei were re-suspended in 200 µl of NIB and mixed with a 140 µl aliquot of melted 2% LMA agarose (Bio-Rad, 1703594) at 43°C, and the mixture was solidified in 50 µl plugs on an ice-cold casting surface. DNA was released by digestion with ESSP (0.1 M EDTA, 1% sodium lauryl sarcosine, 0.2% sodium deoxycholate, 1.48 mg/ml proteinase K) for 36 h at 50°C followed by RNase treatment and extensive washes. High-molecular-weight DNA was stored in TE at 4°C without degradation for up to 8 months.

### Sequencing

The 470-bp (250 paired-end) *Ae. longissima* libraries were sequenced by NovoGene on an Illumina HiSeq 2500. The libraries were sequenced at the Roy J. Carver Biotechnology Center at University of Chicago, Chicago, Illinois. For both *Ae. longissima* and *Ae. speltoides*, 9-kb mate-pair libraries (150 paired-end) were generated and sequenced at the Roy J. Carver Biotechnology Center at UC Illinois. The 10X Genomics chromium libraries (https://www.10xgenomics.com/) were prepared for each genotype following the Chromium Genome library protocol v2 (10X Genomics) and sequenced at the Genome Canada Research and Innovation Centre using the manufacturers’ recommendations across two lanes of Illumina HiSeqX with 150-bp paired-end reads to a minimum 30x coverage. FASTQ files were generated by Longranger (10X Genomics) for analysis (*55*). Hi-C libraries were prepared at the Genome Center at Leibniz Institute for Plant Genetics and Crop Plant Research, Gatersleben, Germany using previously described methods (*56*). Raw data and pseudomolecule sequences were submitted to the European Nucleotide Archive (https://www.ebi.ac.uk/ena) as described in table S5.

### Assembly

The TRITEX pipeline (*30*) was used for genome assembly. Raw data and pseudomolecule sequences were deposited at the European Nucleotide Archive (table S5). Due to high residual heterozygosity in chromosomes 1S and 4S of *Ae. speltoides, de novo* assembly of a Hi-C map did not yield satisfactory results for these two chromosomes, which had to be ordered and oriented solely by collinearity the bread wheat cv. Chinese Spring B-genome (*35*).

### Annotation

Structural gene annotation was done according to the method previously described by (*30*) using *de novo* annotation and homology-based approaches with RNA-seq datasets generated for *Ae. sharonensis* (table S6). Annotation files for the three *Aegilops* genomes are available to download at https://doi.ipk-gatersleben.de:443/DOI/4136d61d-d5a1-4c67-bad1-aa45f7d05dbb/49e438f2-4113-4618-97a6-0c18b6efe6fb/2/1847940088.

Using evidence derived from expression data, RNA-seq data were first mapped using HISAT2 (57) (version 2.0.4, parameter --dta) and subsequently assembled into transcripts by StringTie (58) (version 1.2.3, parameters -m 150-t -f 0.3). *Triticeae* protein sequences from available public datasets (UniProt, 05/10/2016) were aligned against the genome sequence using GenomeThreader (59) (version 1.7.1; arguments -startcodon -finalstopcodon -species rice -gcmincoverage 70 - prseedlength 7 -prhdist 4). All transcripts from RNA-seq and aligned protein sequences were combined using Cuffcompare (*60*) (version 2.2.1) and subsequently merged with Stringtie (version 1.2.3, parameters --merge -m150) into a pool of candidate transcripts. TransDecoder (version 3.0.0) was used to find potential open reading frames and to predict protein sequences within the candidate transcript set.

*Ab initio* annotation using Augustus (*61*) (version 3.3.2) was performed to further improve structural gene annotation. To avoid potential over-prediction, we generated guiding hints using the above RNAseq, protein evidence, and transposable element predictions. A specific model for *Aegilops* was trained using the steps provided in (*62*) and later used for prediction. All structural gene annotations were joined using EvidenceModeller (*63*), and weights were adjusted according to the input source: *ab initio* (2), homology-based (5). Additionally, two rounds of Program to Assemble Spliced Alignments (*64*) (PASA) were run to identify untranslated regions and isoforms using transcripts generated by a genome-guided TRINITY (*65*) (Grabherr 2011) assembly derived from *Ae. sharonensis* RNA-seq data.

We used BLASTP (*66*) (ncbi-blast-2.3.0+, parameters -max_target_seqs 1 -evalue 1e-05) to compare potential protein sequences with a trusted set of reference proteins (Uniprot Magnoliophyta, reviewed/Swissprot, downloaded on 3 Aug 2016). This differentiated candidates into complete and valid genes, non-coding transcripts, pseudogenes, and transposable elements. In addition, we used PTREP (Release 19; http://botserv2.uzh.ch/kelldata/trep-db/index.html), a database of hypothetical proteins containing deduced amino acid sequences in which internal frameshifts have been removed in many cases. This step is particularly useful for the identification of divergent transposable elements with no significant similarity at the DNA level. Best hits were selected for each predicted protein to each of the three databases. Only hits with an E-value below 10e-10 were considered. Furthermore, only hits with subject coverage (for protein references) or query coverage (transposon database) above 95% were considered significant, and protein sequences were further classified using the following confidence: a high-confidence protein sequence was complete and had a subject and query coverage above the threshold in the UniMag database or no BLAST hit in UniMag but in UniPoa and not PTREP; a low-confidence protein sequence was incomplete and had a hit in the UniMag or UniPoa database but not in TREP. Alternatively, it had no hit in UniMag, UniPoa (https://www.uniprot.org), or PTREP, but the protein sequence was complete. The tag REP was assigned for protein sequences not in UniMag and was complete but with hits in PTREP.

Functional annotation of predicted protein sequences was done using the AHRD pipeline {https://github.com/groupschoof/AHRD}. BUSCO (*32*) was used to evaluate the gene space completeness of the pseudomolecule assembly with the ‘embryophyta_odb10’ database containing 1,614 single-copy genes.

### Genome alignments

The *Aegilops* genomes were aligned to CS using Minimap2 (*67*) (https://github.com/lh3/minimap2). The aligned genome was split into 1-kb blocks using the BEDTools (*68*) makewindows command (https://bedtools.readthedocs.io/en/latest/) and then aligned to the CS genome. Results were filtered to include only alignments with MAPQ of 60 and a minimal length of 750 bp.

### NLR annotations

NLR-Annotator (*40*) was used to locate all NLR sequences in the three genomes. A NB-ARC domain global alignment was created through the pipeline from a subset of complete NLRs. Known NLR resistance genes were used as a reference for the tree (table S5). FastTree2 (*69*) (http://www.microbesonline.org/fasttree/) was used to generate a phylogenetic tree from all the NB-ARC sequences. Since the NLR-Annotator pipeline uses NP_001021202.1 (CDP4; Cell Death Protein 4) as an outgroup, we used NP_001021202.1 for tree rooting. We also extracted genes from the whole-genome shotgun sequence annotation by assigned function of ‘Disease’, ‘NBS-LRR’, or ‘NB-ARC’, and these were matched with the NLR-Annotator annotation by position using the intersect option in bedtools (*68*).

Protein sequences of all NLRs were also clustered using CD-HIT (*70*) (https://github.com/weizhongli/cdhit). All the large clusters (n > 30) were located on the tree, and only six large clusters were chosen to represent major branches.

### Phylogeny/Orthologs

OrthoFinder (*36*) (https://github.com/davidemms/OrthoFinder) was used to locate orthologous genes between the *Aegilops* and wheat annotations, and only high-confidence genes were used. We used the three *Aegilops* annotations, *Ae. tauschii*, the AB annotation of WEW, and ABD annotation of CS. The polyploid annotations (CS and WEW) were split into the subgenomes, and each was handled separately (table S6). Additionally, OrthoFinder was used to identify orthologous NLR sequences (that were a subset from the gene annotation) and to analyze the phylogenetic relationship between the genomes and subgenomes. To compare the gene annotations, BLASTN was used to align the high-confidence genes from *Ae. longissima* and *Ae. sharonensis* to each other and to the B and D subgenomes of CS and high-confidence genes from *Ae. speltoides* to the B subgenomes of CS and WEW.

### Haploblocks

Haploblock analysis was performed using the methods described by (*37*). Modifications were made in terms of a lowered percent identity for some pair-wise comparisons to reflect the phylogenetic distance.

## Acknowledgments

We are grateful to the Harold and Adele Lieberman Germplasm Bank, Tel Aviv University, for making *Ae. longissima, Ae. sharonensis*, and WEW seeds available, and to the Lieberman Family and JNF Australia for the generous support of this project. We kindly acknowledge Manuela Knauft and Ines Walde for technical assistance on Hi-C library preparation and sequencing. We thank Prof. Moshe Feldman for helpful discussions on the section Sitopsis. We thank Anne Fiebig for sequence data submission.

## Author contributions

Extracted DNA for whole-genome shotgun sequencing and long mate-pair libraries: AM, JD, HS; Performed 10X Chromium sequencing: JE, CJP; Performed Hi-C sequencing: AH, NS; Contribution of the *Ae. sharonensis* genome: GY, BW; Prepared material and involved in *Ae. speltoides* sequencing: HS, AD, CJP; Assembled the *Ae. longissima* and *A. speltoides* genomes: MM, RA; Gene Annotation: TL, HG, MS; Performed bioinformatics analyses including whole genome alignment, haploblocks, OrthoFinder and BUSCO: TL, MS, RA. Performed NLR analysis: RA, TL, BS, HS; Provided scientific support: EM, KM. Conceived study: AS, RA, BW, HS; drafted manuscript: RA, AS, BW. All co-authors read and approved the final manuscript. Authors are grouped by institution in the author list, except for the first four and last four authors.

## Data and materials availability

The datasets generated during and/or analyzed during the current study are publicly available as follows. The sequence reads and the genome assembly were deposited in the European Nucleotide Archive under project number PRJEB41661, PRJEB41746, PRJEB40543, PRJEB40544, PRJEB40050, and PRJEB40051, respectively. Gene annotation files are available at: https://doi.ipk-gatersleben.de/DOI/4136d61d-d5a1-4c67-bad1-aa45f7d05dbb/49e438f2-4113-4618-97a6-0c18b6efe6fb/2/1847940088. Seeds of *Ae. longissima* accession AEG-6782-2 and *Ae. speltoides* accession AEG-9674-1 are available from the Institute for Cereal Crops Improvement at Tel Aviv University (https://en-lifesci.tau.ac.il/icci).

## Supplementary Materials

Supplementry.pdf

## Supplementary Materials for

**Fig. S1.**
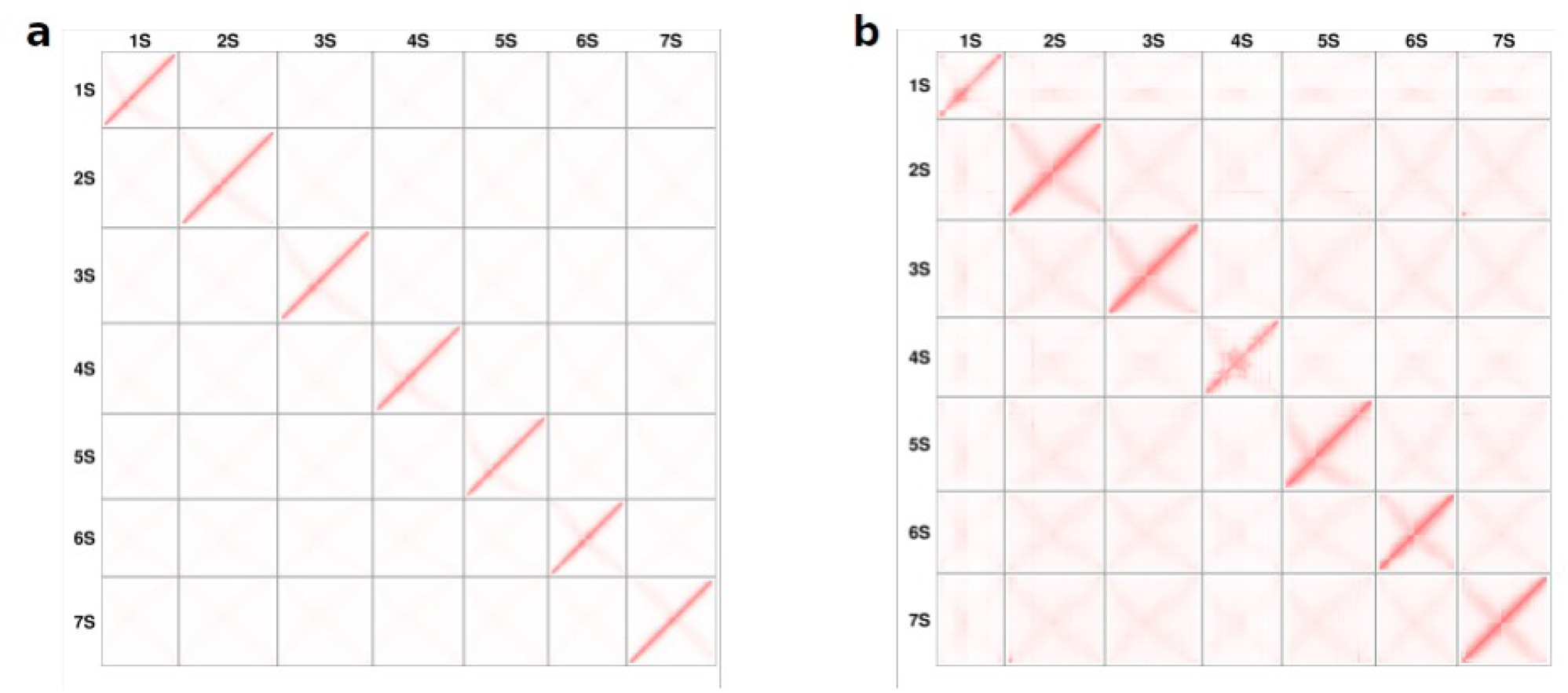
Hi-C contact matrices for **a**, *Ae. longissima* and **b**, *Ae. speltoides*.

**Fig. S2.**
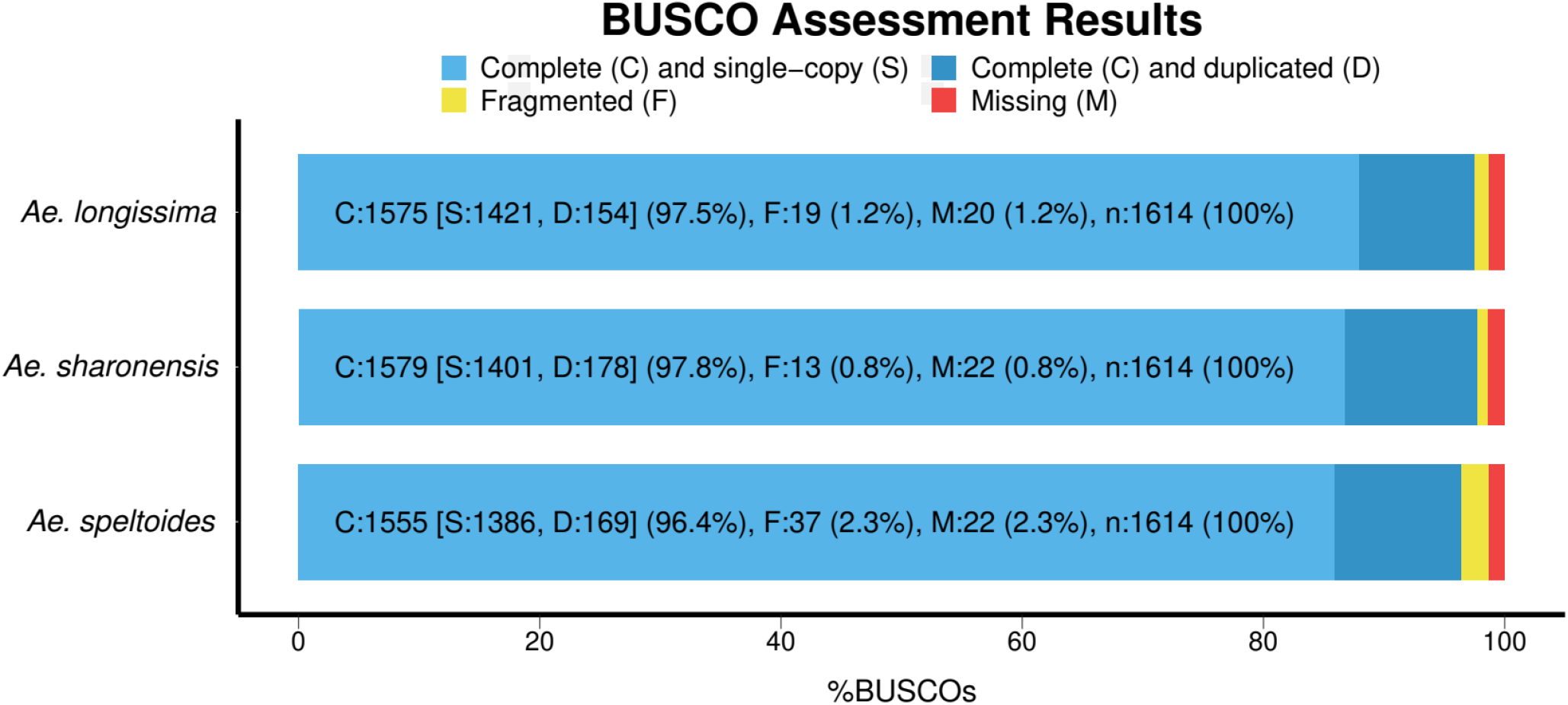
BUSCO assessment of the completeness of the gene annotation using the ‘embryophyta_odb10’ database that includes 1,614 core plant genes (*32*).

**Fig. S3.**
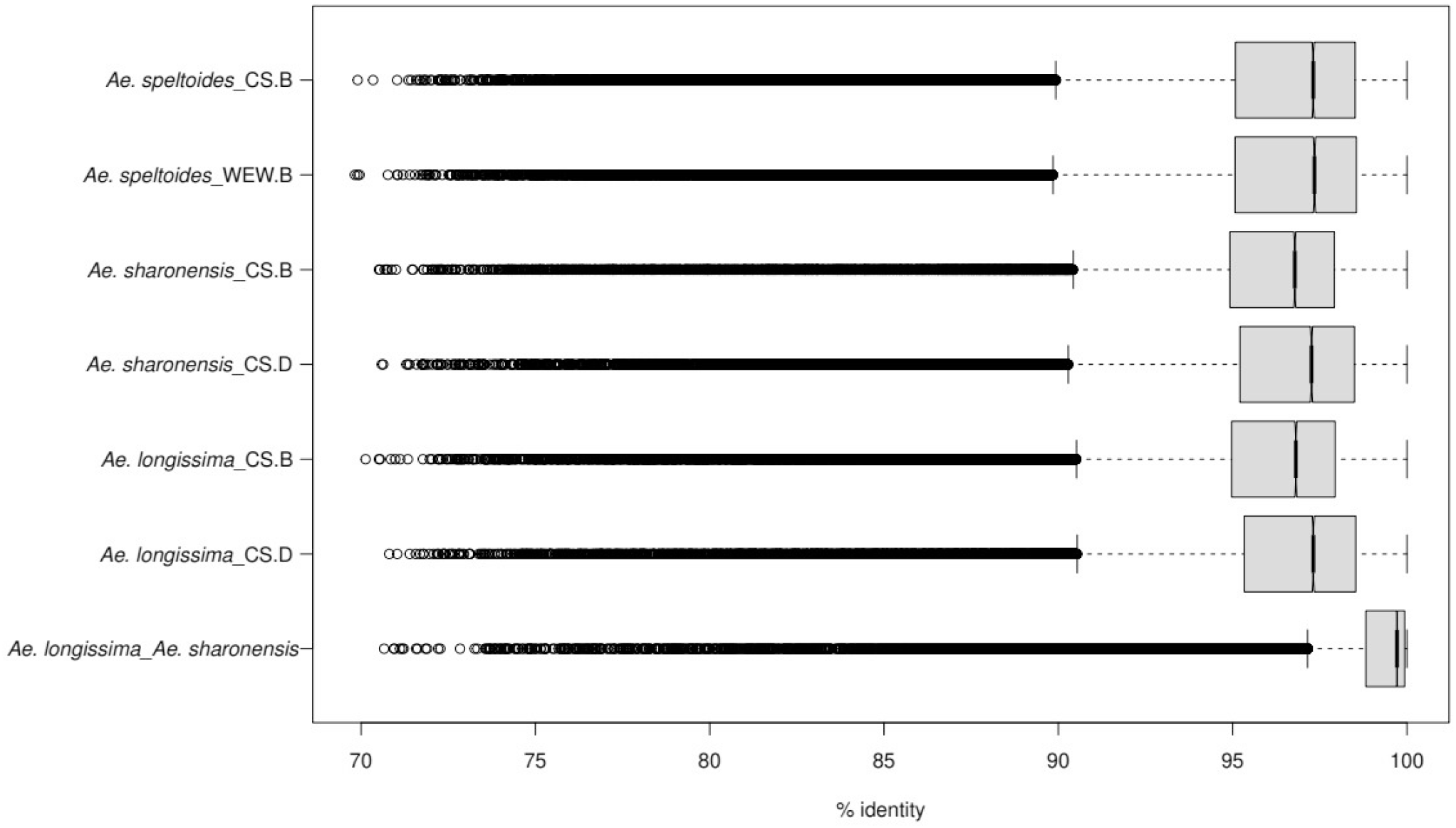
Gene annotation pair alignments. Box plots show the percentage of identity between the best hit for each query (first label name) gene in the reference (second label name) annotation.

**Fig. S4.** This figure is included as a separate PDF file. Detailed phylogenetic tree of NLR genes in *Aegilops* and wheat. Cloned genes (Supplementary Table 5) were added to the tree, and their branches are highlighted. Tip labels refer to NLR-Annotator IDs. Tip point color shows whether a match is found between the NLR annotation (NLR-Annotator) and the whole-genome annotation; gray point means no match. WEW, wild emmer wheat, *Triticum turgidum* ssp. *dicoccoides*. CS, bread wheat, *Triticum aestivum* cv. Chinese Spring.

**Fig. S5.**
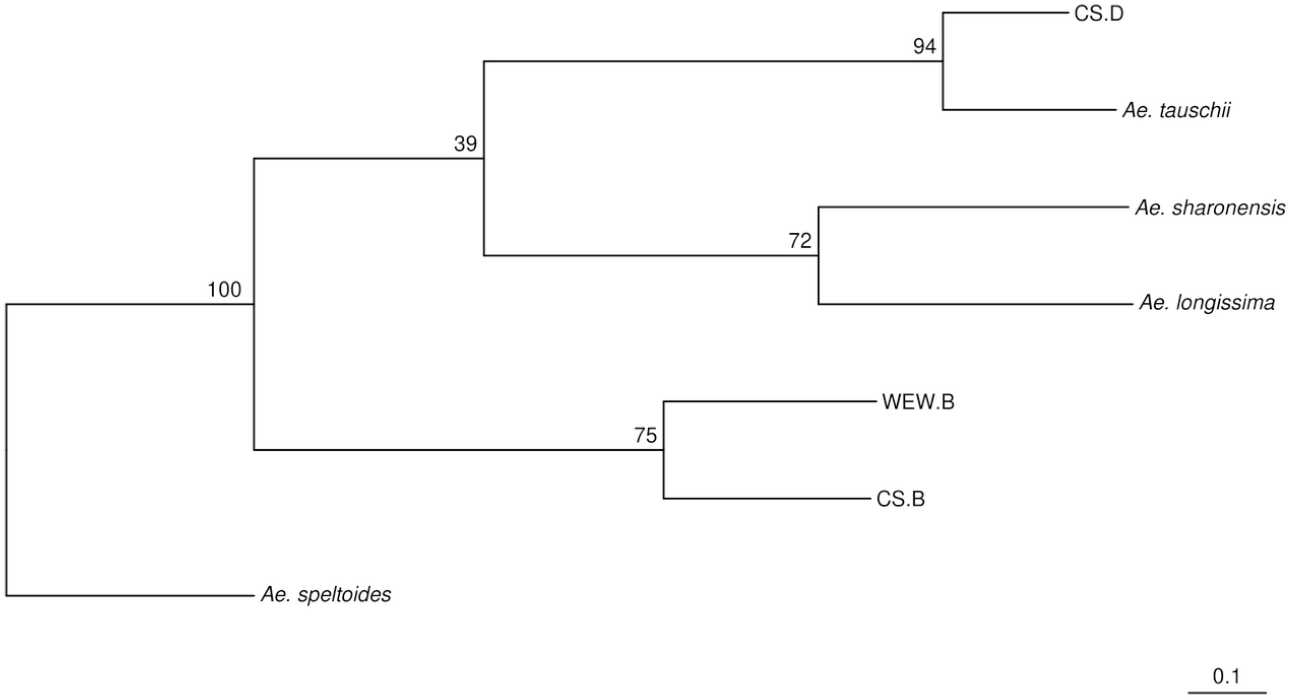
Phylogenetic tree generated from the OrthoFinder analysis with all NLR gene predictions as input. Values correspond to OrthoFinder-based support values.

**Table S1.** Overview of the three *Aegilops* assemblies showing all chromosomes including chromosome “Un” with all unassociated scaffolds.

| Chromosome | Chromosome length (bp) | <sup>a</sup>No. HC genes | <sup>b</sup>No. LC genes | Total No. genes | N50 (bp) | N90 (bp) | #scaffolds |
| --- | --- | --- | --- | --- | --- | --- | --- |
| 1S | 742,303,966 | 3,438 | 3,973 | 7,411 | 4,561,840 | 1,138,021 | 289 |
| 2S | 956,173,857 | 4,644 | 5,107 | 9,751 | 4,849,770 | 1,135,822 | 362 |
| 3S | 916,702,776 | 4,531 | 4,957 | 9,488 | 4,807,476 | 999,978 | 368 |
| 4S | 874,517,040 | 3,697 | 4,426 | 8,123 | 4,768,090 | 1,074,529 | 346 |
| 5S | 816,294,110 | 4,328 | 4,595 | 8,923 | 4,330,990 | 1,052,266 | 347 |
| 6S | 750,216,944 | 3,259 | 4,082 | 7,341 | 3,936,747 | 1,048,299 | 312 |
| 7S | 862,608,691 | 3,680 | 4,814 | 8,494 | 4,285,042 | 1,113,450 | 339 |
| Un | 783,655,401 | 3,606 | 4,956 | 8,562 | 56,180 | 1,491 | 127,984 |
| Total | 6,702,472,785 | 31,183 | 36,910 | 68,093 | 3,754,329 | 242,603 | 130,347 |

*Ae. speltoides*
| Chromosome | Chromosome length (bp) | <sup>a</sup>No. HC genes | <sup>b</sup>No. LC genes | Total No. genes | N50 (bp) | N90 (bp) | #scaffolds |
| --- | --- | --- | --- | --- | --- | --- | --- |
| 1S | 445,829,560 | 3,178 | 4,484 | 7,662 | 1,338,029 | 378,510 | 529 |
| 2S | 657,893,865 | 5,522 | 6,750 | 12,272 | 7,145,516 | 1,121,567 | 281 |
| 3S | 636,117,214 | 5,137 | 6,359 | 11,496 | 14,542,279 | 1,329,581 | 209 |
| 4S | 520,569,408 | 3,149 | 4,127 | 7,276 | 2,294,611 | 639,716 | 386 |
| 5S | 614,738,994 | 5,113 | 6,471 | 11,584 | 9,396,929 | 1,445,106 | 203 |
| 6S | 536,175,046 | 3,861 | 5,350 | 9,211 | 6,487,485 | 1,624,515 | 188 |
| 7S | 610,578,938 | 4,538 | 6,477 | 11,015 | 7,894,947 | 1,022,973 | 264 |
| Un | 1,114,993,717 | 6,430 | 18,109 | 24,539 | 21,791 | 1,622 | 185,157 |
| Total | 5,136,896,742 | 36,928 | 58,127 | 95,055 | 3,111,390 | 20,000 | 187,217 |

*Ae. sharonensis*<sup>c</sup>
| Chromosome<sup>c</sup> | Chromosome length (bp)<sup>c</sup> | <sup>a</sup>No. HC genes | <sup>b</sup>No. LC genes | Total No. genes | N50 (bp)<sup>c</sup> | N90 (bp)<sup>c</sup> | #scaffolds<sup>c</sup> |
| --- | --- | --- | --- | --- | --- | --- | --- |
| 1S | 782,818,162 | 3,678 | 4,396 | 8,074 | 14,301,836 | 1,946,926 | 168 |
| 2S | 1,022,071,454 | 5,093 | 5,760 | 10,853 | 12,458,785 | 2,598,021 | 182 |
| 3S | 971,920,087 | 4,983 | 5,579 | 10,562 | 17,681,665 | 2,964,762 | 161 |
| 4S | 827,198,496 | 3,298 | 4,016 | 7,314 | 17,868,812 | 2,716,220 | 138 |
| 5S | 867,619,200 | 4,713 | 4,961 | 9,674 | 13,883,202 | 3,670,029 | 126 |
| 6S | 806,566,123 | 3,599 | 4,568 | 8,167 | 13,434,032 | 2,801,261 | 123 |
| 7S | 1,015,700,474 | 4,710 | 6,119 | 10,829 | 9,208,414 | 1,394,866 | 268 |
| Un | 420,139,911 | 1,124 | 2,200 | 3,324 | 15,399 | 1,188 | 93,480 |
| Total | 6,714,033,907 | 31,198 | 37,599 | 68,797 | 12,378,845 | 1,132,218 | 94,646 |

<sup>a</sup>HC, high-confidence genes.<sup>b</sup>LC, low-confidence genes.<sup>c</sup>Adapted from<sup>29</sup>

**Table S2.** Summary of OrthoFinder results for all high-confidence genes.

|  | Number of genes | Number of genes in orthogroups | Number of unassigned genes | % of genes in orthogroups | % of unassigned genes | Number of orthogroups containing species | % of orthogroups containing species | Number of species-specific orthogroups | Number of genes in species-specific orthogroups | % of genes in species-specific orthogroups |
| --- | --- | --- | --- | --- | --- | --- | --- | --- | --- | --- |
| *Ae. longissima* | 31,183 | 30,902 | 281 | 99.1 | 0.9 | 23,633 | 72 | 15 | 102 | 0.3 |
| *Ae. sharonensis* | 31,198 | 30,961 | 237 | 99.2 | 0.8 | 23,559 | 71.8 | 14 | 40 | 0.1 |
| *Ae. speltoides* | 36,928 | 36,327 | 601 | 98.4 | 1.6 | 24,427 | 74.4 | 136 | 1,083 | 2.9 |
| WEW.A | 32,706 | 31,943 | 763 | 97.7 | 2.3 | 25,222 | 76.9 | 69 | 158 | 0.5 |
| WEW.B | 33,052 | 32,222 | 830 | 97.5 | 2.5 | 24,679 | 75.2 | 129 | 353 | 1.1 |
| CS.A | 35,275 | 34,656 | 619 | 98.2 | 1.8 | 26,354 | 80.3 | 58 | 132 | 0.4 |
| CS.B | 35,561 | 34,903 | 658 | 98.1 | 1.9 | 25,706 | 78.3 | 46 | 101 | 0.3 |
| CS.D | 34,161 | 33,627 | 534 | 98.4 | 1.6 | 25,813 | 78.7 | 34 | 79 | 0.2 |
| *Ae. tauschii* | 39,622 | 37,318 | 2304 | 94.2 | 5.8 | 25,720 | 78.4 | 510 | 2,398 | 6.1 |

WEW, wild emmer wheat, *Triticum turgidum* ssp. *dicoccoides*.
CS, bread wheat, *Triticum aestivum* cv. Chinese Spring.

**Table S3.** Number of predicted NLR genes in different genomes.

| Species | <sup>a</sup>NLR-Annotator | <sup>b</sup>PGSB annotation |
| --- | --- | --- |
| *Ae. sharonensis* | 1,451 (979) | 1,143 (773) |
| *Ae. longissima* | 1,555 (978) | 1,065 (732) |
| *Ae. speltoides* | 1,358 (721) | 1,570 (1,045) |
| *Ae. tauschii* | 888 |  |
| *T. aestivum* (CS) | 3,184 (A, 1,064; B, 1,198; D, 922) |  |
| *T. dicoccoides* (WEW) | 2,230 (A, 1,015; B, 1,215) |  |

<sup>a</sup>Number represents all predicted genes. The number in parentheses represents only complete genes. <sup>b</sup>Number represents disease resistance + NBS-NLR + NB-ARC genes. The number in parentheses represents only NBS-NLR + NB-ARC genes. WEW, wild emmer wheat, *Triticum turgidum* ssp. *dicoccoides*. CS, bread wheat, *Triticum aestivum* cv. Chinese Spring.

**Table S4.**
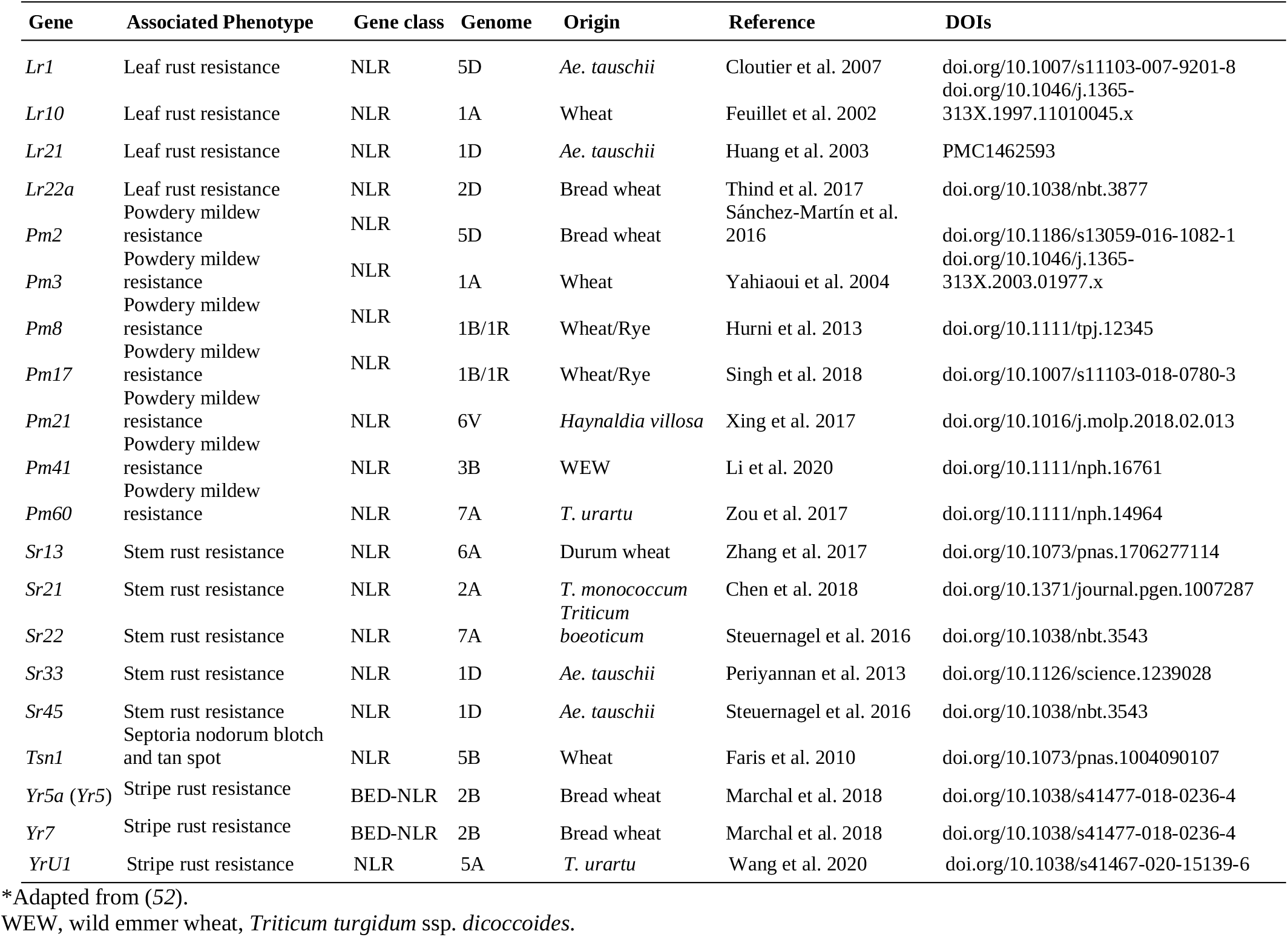
Cloned NLR genes* used as reference for the NLR phylogenetic tree.

**Table S5.**
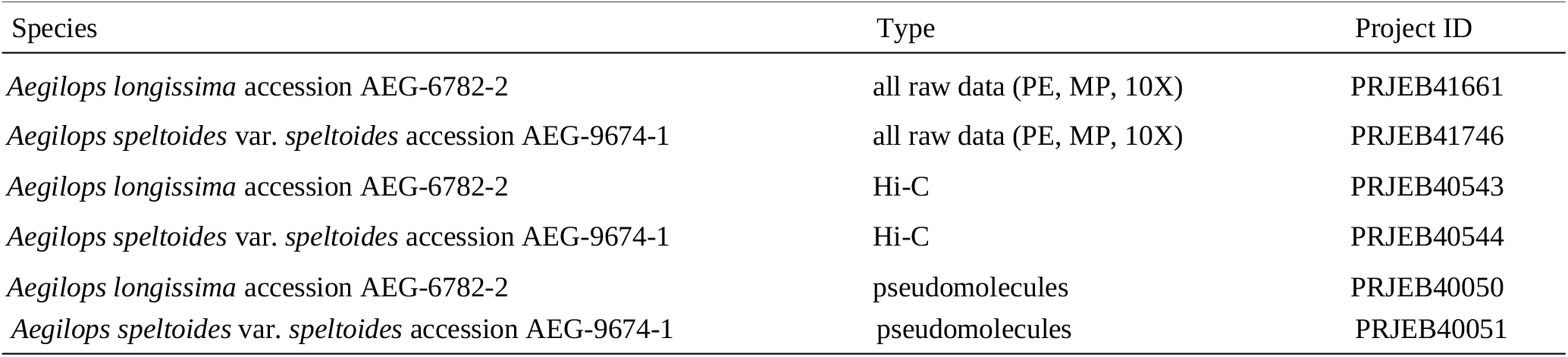
Project numbers for European Nucleotide Archive raw sequencing data and pseudomolecule submission.

**Table S6.**
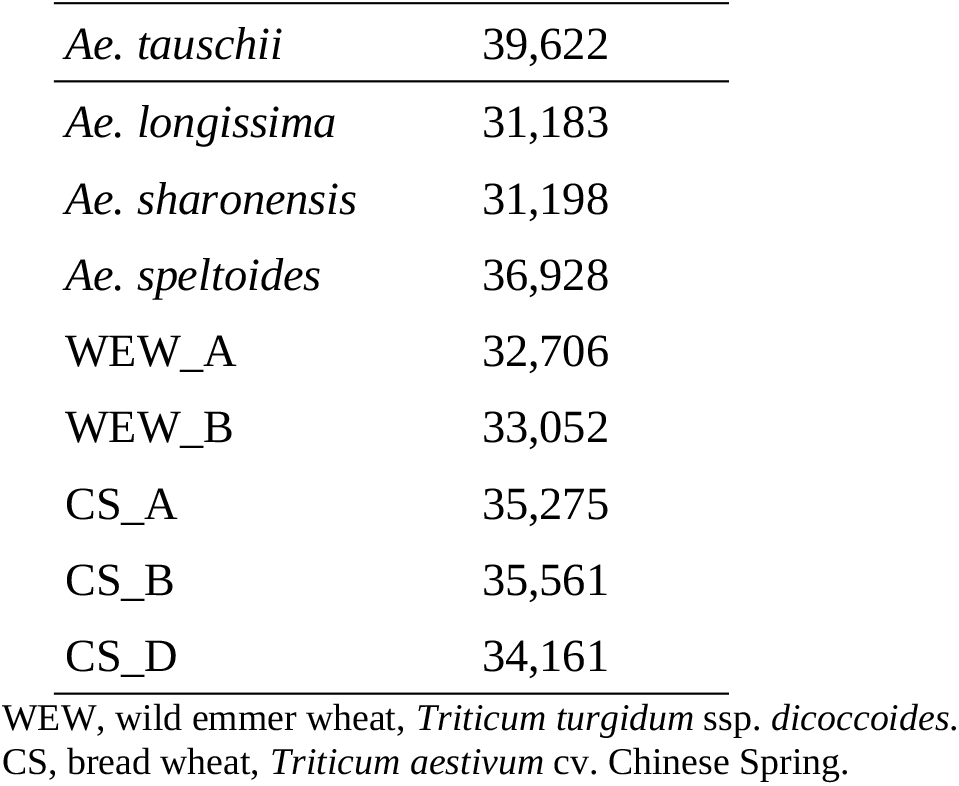
Number of high-confidence genes per species used for ortholog phylogenetic analysis.

## Data S1. (separate file)

dataset_1.xlsx

Summary tables of haploblocks analysis

## References

1. D. Zohary, M. Hopf, E. Weiss, Domestication of Plants in the Old World: The origin and spread of domesticated plants in Southwest Asia, Europe, and the Mediterranean Basin. Domest. Plants Old World Orig. Spread Domest. Plants Southwest Asia, Eur. Mediterr. Basin, 1–264 (2012).

2. R. J. Giles, T. A. Brown, GluDy allele variations in Aegilops tauschii and Triticum aestivum: Implications for the origins of hexaploid wheats. Theor. Appl. Genet. 112, 1563–1572 (2006).

3. A. M. Mastrangelo, L. Cattivelli, What Makes Bread and Durum Wheat Different? Trends Plant Sci. (2021).

4. J. Dubcovsky, J. Dvorak, Genome plasticity a key factor in the success of polyploid wheat under domestication. Science. 316, 1862–1866 (2007).

5. J. C. Reif, P. Zhang, S. Dreisigacker, M. L. Warburton, M. Van Ginkel, D. Hoisington, M. Bohn, A. E. Melchinger, Wheat genetic diversity trends during domestication and breeding. Theor. Appl. Genet. 110, 859–864 (2005).

6. C. Pont, T. Leroy, M. Seidel, A. Tondelli, W. Duchemin, D. Armisen, D. Lang, D. Bustos-Korts, N. Goué, F. Balfourier, M. Molnár-Láng, J. Lage, B. Kilian, H. Özkan, D. Waite, S. Dyer, T. Letellier, M. Alaux, J. Russell, B. Keller, F. van Eeuwijk, M. Spannagl, K. F. X. Mayer, R. Waugh, N. Stein, L. Cattivelli, G. Haberer, G. Charmet, J. Salse, C. Saintenac, P. Lasserre-Zuber, M. R. Perretant, A. Didier, S. Bouchet, J. Boudet, E. Bancel, M. Merlino, C. Grand-Ravel, T. Langin, M. Bayer, A. Booth, I. Dawson, P. Schweizer, K. Neumann, G. Kema, M. Bink, M. Molnar-Lang, M. Megyeri, P. Miko, G. Linc, J. Wright, L. Clissold, K. Krasileva, J. De Vega, P. Bailey, V. Goody, S. Wilbraham, M. Anissi, J. Moore, D. Swan, C. Watkins, D. L. M. Spannagl, A. Korol, T. Krugman, T. Fahima, L. Rossini, H. Jones, N. Morris, A. Costanzo, T. Wicker, T. Muller, M. Martelli, S. Ravaglia, C. Bonard, S. Crépieux, J. Saranga, E. Çakır, Tracing the ancestry of modern bread wheats. Nat. Genet. 51, 905–911 (2019).

7. R. A. Finch, T. E. Miller, M. D. Bennett, “Cuckoo” Aegilops addition chromosome in wheat ensures its transmission by causing chromosome breaks in meiospores lacking it. Chromosoma. 90, 84–88 (1984).

8. H. Tsujimoto, Two new sources of gametocidal genes from Ae. longissima and Ae. sharonensis. Wheat Inf. Serv. 79, 42–46 (1994).

9. B. Kilian, K. Mammen, E. Millet, R. Sharma, A. Graner, F. Salamini, K. Hammer, H. Özkan, Aegilops. Wild Crop Relat. Genomic Breed. Resour., 1–76 (2011).

10. S. Khazan, A. Minz-Dub, H. Sela, J. Manisterski, P. Ben-Yehuda, A. Sharon, E. Millet, Reducing the size of an alien segment carrying leaf rust and stripe rust resistance in wheat. BMC Plant Biol. 20 (2020).

11. S. Arora, B. Steuernagel, K. Gaurav, S. Chandramohan, Y. Long, O. Matny, R. Johnson, J. Enk, S. Periyannan, N. Singh, M. Asyraf Md Hatta, N. Athiyannan, J. Cheema, G. Yu, N. Kangara, S. Ghosh, L. J. Szabo, J. Poland, H. Bariana, J. D. G. Jones, A. R. Bentley, M. Ayliffe, E. Olson, S. S. Xu, B. J. Steffenson, E. Lagudah, B. B. H. Wulff, Resistance gene cloning from a wild crop relative by sequence capture and association genetics. Nat. Biotechnol. 37, 139–143 (2019).

12. C. Uauy, B. B. H. Wulff, J. Dubcovsky, Combining Traditional Mutagenesis with New High-Throughput Sequencing and Genome Editing to Reveal Hidden Variation in Polyploid Wheat. Annu. Rev. Genet. 51, 435–454 (2017).

13. J. Wang, M. C. Luo, Z. Chen, F. M. You, Y. Wei, Y. Zheng, J. Dvorak, Aegilops tauschii single nucleotide polymorphisms shed light on the origins of wheat D-genome genetic diversity and pinpoint the geographic origin of hexaploid wheat. New Phytol. 198, 925–937 (2013).

14. M. C. Luo, Y. Q. Gu, D. Puiu, H. Wang, S. O. Twardziok, K. R. Deal, N. Huo, T. Zhu, L. Wang, Y. Wang, P. E. McGuire, S. Liu, H. Long, R. K. Ramasamy, J. C. Rodriguez, L. Van Sonny, L. Yuan, Z. Wang, Z. Xia, L. Xiao, O. D. Anderson, S. Ouyang, Y. Liang, A. V. Zimin, G. Pertea, P. Qi, J. L. Bennetzen, X. Dai, M. W. Dawson, H. G. Müller, K. Kugler, L. Rivarola-Duarte, M. Spannagl, K. F. X. Mayer, F. H. Lu, M. W. Bevan, P. Leroy, P. Li, F. M. You, Q. Sun, Z. Liu, E. Lyons, T. Wicker, S. L. Salzberg, K. M. Devos, J. Dvoák, Genome sequence of the progenitor of the wheat D genome Aegilops tauschii. Nature. 551, 498–502 (2017).

15. M. Kishii, An update of recent use of Aegilops species in wheat breeding. Front. Plant Sci. 10 (2019).

16. K. Kerby, J. Kuspira, The phylogeny of the polyploid wheats Triticum aestivum (bread wheat) and Triticum turgidum (macaroni wheat). Genome. 29, 722–737 (1987).

17. T. Marcussen, S. R. Sandve, L. Heier, M. Spannagl, M. Pfeifer, K. S. Jakobsen, B. B. H. Wulff, B. Steuernagel, K. F. X. Mayer, O. A. Olsen, J. Rogers, J. Doleželz, C. Pozniak, K. Eversole, C. Feuillet, B. Gill, B. Friebe, A. J. Lukaszewski, P. Sourdille, T. R. Endo, M. Kubaláková, J. Šíhalíková, Z. Dubská, J. Vrána, R. Šperková, H. Šimková, M. Febrer, L. Clissold, K. McLay, K. Singh, P. Chhuneja, N. K. Singh, J. Khurana, E. Akhunov, F. Choulet, A. Alberti, V. Barbe, P. Wincker, H. Kanamori, F. Kobayashi, T. Itoh, T. Matsumoto, H. Sakai, T. Tanaka, J. Wu, Y. Ogihara, H. Handa, P. R. Maclachlan, A. Sharpe, D. Klassen, D. Edwards, J. Batley, S. R. Sandve, S. Lien, B. Wulff, M. Caccamo, S. Ayling, R. H. Ramirez-Gonzalez, B. J. Clavijo, J. Wright, M. M. Martis, M. Mascher, J. Chapman, J. A. Poland, U. Scholz, K. Barry, R. Waugh, D. S. Rokhsar, G. J. Muehlbauer, N. Stein, H. Gundlach, M. Zytnicki, V. Jamilloux, H. Quesneville, T. Wicker, P. Faccioli, M. Colaiacovo, A. M. Stanca, H. Budak, L. Cattivelli, N. Glover, L. Pingault, E. Paux, S. Sharma, R. Appels, M. Bellgard, B. Chapman, T. Nussbaumer, K. C. Bader, H. Rimbert, S. Wang, R. Knox, A. Kilian, M. Alaux, F. Alfama, L. Couderc, N. Guilhot, C. Viseux, M. Loaec, B. Keller, S. Praud, Ancient hybridizations among the ancestral genomes of bread wheat. Science. 345 (2014).

18. G. Petersen, O. Seberg, M. Yde, K. Berthelsen, Phylogenetic relationships of Triticum and Aegilops and evidence for the origin of the A, B, and D genomes of common wheat (Triticum aestivum). Mol. Phylogenet. Evol. 39, 70–82 (2006).

19. K. Yamane, T. Kawahara, Intra-and interspecific phylogenetic relationships among diploid Triticum-aegilops species (Poaceae) based on base-pair substitutions, indels, and microsatellites in chloroplast noncoding sequences. Am. J. Bot. 92, 1887–1898 (2005).

20. The International Wheat Genome Sequencing Consortium, A chromosome-based draft sequence of the hexaploid bread wheat (Triticum aestivum) genome. Science. 345, 1251788 (2014).

21. N. Bernhardt, J. Brassac, X. Dong, E. M. Willing, C. H. Poskar, B. Kilian, F. R. Blattner, Genome-wide sequence information reveals recurrent hybridization among diploid wheat wild relatives. Plant J. 102, 493–506 (2020).

22. O. U. Edet, Y. S. A. Gorafi, S. Nasuda, H. Tsujimoto, DArTseq-based analysis of genomic relationships among species of tribe Triticeae. Sci. Rep. 8 (2018)

23. S. Glémin, C. Scornavacca, J. Dainat, C. Burgarella, V. Viader, M. Ardisson, G. Sarah, S. Santoni, J. David, V. Ranwez, Pervasive hybridizations in the history of wheat relatives. Sci. Adv. 5 (2019).

24. S. Huynh, T. Marcussen, F. Felber, C. Parisod, Hybridization preceded radiation in diploid wheats. Mol. Phylogenet. Evol. 139, 599068 (2019).

25. Y. Anikster, J. Manisterski, D. L. Long, K. J. Leonard, Resistance to leaf rust, stripe rust, and stem rust in Aegilops spp. in Israel. Plant Dis. 89, 303–308 (2005).

26. S. Huang, B. J. Steffenson, H. Sela, K. Stinebaugh, Resistance of Aegilops longissima to the rusts of wheat. Plant Dis. 102, 1124–1135 (2018).

27. P. D. Olivera, J. A. Kolmer, Y. Anikster, B. J. Steffenson, Resistance of Sharon goatgrass (Aegilops sharonensis) to fungal diseases of wheat. Plant Dis. 91, 942–950 (2007).

28. J. C. Scott, J. Manisterski, H. Sela, P. Ben-Yehuda, B. J. Steffenson, Resistance of Aegilops species from israel to widely virulent african and israeli races of the wheat stem rust pathogen. Plant Dis. 98, 1309–1320 (2014).

29. G. Yu, O. Matny, N. Champouret, B. Steuernagel, M. J. Moscou, I. Hernández-Pinzón, P. Green, S. Hayta, M. Smedley, W. Harwood, N. Kangara, Y. Yue, C. Gardener, M. J. Banfield, P. D. Olivera, C. Welchin, J. Simmons, E. Millet, A. Minz-Dub, M. Ronen, R. Avni, A. Sharon, M. Patpour, A. F. Justesen, M. Jayakodi, A. Himmelbach, N. Stein, S. Wu, J. Poland, J. Ens, C. Pozniak, M. Karafiátová, I. Molnár, J. Doležel, E. R. Ward, T. L. Reuber, J. D. G. Jones, M. Mascher, B. J. Steffenson, B. B. H. Wulff, Running title: Reference genome-assisted identification of the stem rust resistance gene. Nat. Commun. (In review) (2021).

30. C. Monat, S. Padmarasu, T. Lux, T. Wicker, H. Gundlach, A. Himmelbach, J. Ens, C. Li, G. J. Muehlbauer, A. H. Schulman, R. Waugh, I. Braumann, C. Pozniak, U. Scholz, K. F. X. Mayer, M. Spannagl, N. Stein, M. Mascher, TRITEX: Chromosome-scale sequence assembly of Triticeae genomes with open-source tools. Genome Biol. 20 (2019).

31. T. Eilam, Y. Anikster, E. Millet, J. Manisterski, M. Feldman, Nuclear DNA amount and genome downsizing in natural and synthetic allopolyploids of the genera Aegilops and Triticum. Genome. 51, 616–627 (2008).

32. F. A. Simão, R. M. Waterhouse, P. Ioannidis, E. V. Kriventseva, E. M. Zdobnov, BUSCO: Assessing genome assembly and annotation completeness with single-copy orthologs. Bioinformatics. 31, 3210–3212 (2015).

33. H. Zhang, S. M. Reader, X. Liu, J. Z. Jia, M. D. Gale, K. M. Devos, Comparative genetic analysis of the Aegilops longissima and Ae. sharonensis genomes with common wheat. Theor. Appl. Genet. 103, 518–525 (2001).

34. R. Avni, M. Nave, O. Barad, K. Baruch, S. O. Twardziok, H. Gundlach, I. Hale, M. Mascher, M. Spannagl, K. Wiebe, K. W. Jordan, G. Golan, J. Deek, B. Ben-Zvi, G. Ben-Zvi, A. Himmelbach, R. P. Maclachlan, A. G. Sharpe, A. Fritz, R. Ben-David, H. Budak, T. Fahima, A. Korol, J. D. Faris, A. Hernandez, M. A. Mikel, A. A. Levy, B. Steffenson, M. Maccaferri, R. Tuberosa, L. Cattivelli, P. Faccioli, A. Ceriotti, K. Kashkush, M. Pourkheirandish, T. Komatsuda, T. Eilam, H. Sela, A. Sharon, N. Ohad, D. A. Chamovitz, K. F. X. Mayer, N. Stein, G. Ronen, Z. Peleg, C. J. Pozniak, E. D. Akhunov, A. Distelfeld, Wild emmer genome architecture and diversity elucidate wheat evolution and domestication. Science. 357, 93–97 (2017).

35. R. Appels, K. Eversole, C. Feuillet, B. Keller, J. Rogers, N. Stein, C. J. Pozniak, F. Choulet, A. Distelfeld, J. Poland, G. Ronen, O. Barad, K. Baruch, G. Keeble-Gagnère, M. Mascher, G. Ben-Zvi, A. A. Josselin, A. Himmelbach, F. Balfourier, J. Gutierrez-Gonzalez, M. Hayden, C. S. Koh, G. Muehlbauer, R. K. Pasam, E. Paux, P. Rigault, J. Tibbits, V. Tiwari, M. Spannagl, D. Lang, H. Gundlach, G. Haberer, K. F. X. Mayer, D. Ormanbekova, V. Prade, T. Wicker, D. Swarbreck, H. Rimbert, M. Felder, N. Guilhot, G. Kaithakottil, J. Keilwagen, P. Leroy, T. Lux, S. Twardziok, L. Venturini, A. Juhasz, M. Abrouk, I. Fischer, C. Uauy, P. Borrill, R. H. Ramirez-Gonzalez, D. Arnaud, S. Chalabi, B. Chalhoub, A. Cory, R. Datla, M. W. Davey, J. Jacobs, S. J. Robinson, B. Steuernagel, F. Van Ex, B. B. H. Wulff, M. Benhamed, A. Bendahmane, L. Concia, D. Latrasse, M. Alaux, J. Bartoš, A. Bellec, H. Berges, J. Doležel, Z. Frenkel, B. Gill, A. Korol, T. Letellier, O. A. Olsen, H. Šimková, K. Singh, M. Valárik, E. Van Der Vossen, S. Vautrin, S. Weining, T. Fahima, V. Glikson, D. Raats, H. Toegelová, J. Vrána, P. Sourdille, B. Darrier, D. Barabaschi, L. Cattivelli, P. Hernandez, S. Galvez, H. Budak, J. D. G. Jones, K. Witek, G. Yu, I. Small, J. Melonek, R. Zhou, T. Belova, K. Kanyuka, R. King, K. Nilsen, S. Walkowiak, R. Cuthbert, R. Knox, K. Wiebe, D. Xiang, A. Rohde, T. Golds, J. Čížkova, B. A. Akpinar, S. Biyiklioglu, L. Gao, A. N’Daiye, J. Číhalíková, M. Kubaláková, J. Šafář, F. Alfama, A. F. Adam-Blondon, R. Flores, C. Guerche, M. Loaec, H. Quesneville, A. G. Sharpe, J. Condie, J. Ens, R. Maclachlan, Y. Tan, A. Alberti, J. M. Aury, V. Barbe, A. Couloux, C. Cruaud, K. Labadie, S. Mangenot, P. Wincker, G. Kaur, M. Luo, S. Sehgal, P. Chhuneja, O. P. Gupta, S. Jindal, P. Kaur, P. Malik, P. Sharma, B. Yadav, N. K. Singh, J. P. Khurana, C. Chaudhary, P. Khurana, V. Kumar, A. Mahato, S. Mathur, A. Sevanthi, N. Sharma, R. S. Tomar, K. Holušová, O. Plíhal, M. D. Clark, D. Heavens, G. Kettleborough, J. Wright, B. Balcárková, Y. Hu, N. Ravin, K. Skryabin, A. Beletsky, V. Kadnikov, A. Mardanov, M. Nesterov, A. Rakitin, E. Sergeeva, H. Kanamori, S. Katagiri, F. Kobayashi, S. Nasuda, T. Tanaka, J. Wu, F. Cattonaro, M. Jiumeng, K. Kugler, M. Pfeifer, S. Sandve, X. Xun, B. Zhan, J. Batley, P. E. Bayer, D. Edwards, S. Hayashi, Z. Tulpová, P. Visendi, L. Cui, X. Du, K. Feng, X. Nie, W. Tong, L. Wang, Shifting the limits in wheat research and breeding using a fully annotated reference genome. Science. 361 (2018).

36. D. M. Emms, S. Kelly, OrthoFinder: Phylogenetic orthology inference for comparative genomics. Genome Biol. 20 (2019).

37. J. Brinton, R. H. Ramirez-Gonzalez, J. Simmonds, L. Wingen, S. Orford, S. Griffiths, G. Haberer, M. Spannagl, S. Walkowiak, C. Pozniak, C. Uauy, A haplotype-led approach to increase the precision of wheat breeding. Commun. Biol. 3 (2020).

38. A. L. Delcher, A. Phillippy, J. Carlton, S. L. Salzberg, Fast algorithms for large-scale genome alignment and comparison. Nucleic Acids Res. 30, 2478–2483 (2002).

39. J. Kourelis, R. A. L. Van Der Hoorn, Defended to the nines: 25 years of resistance gene cloning identifies nine mechanisms for R protein function. Plant Cell. 30, 285–299 (2018).

40. B. Steuernagel, K. Witek, S. G. Krattinger, R. H. Ramirez-Gonzalez, H. J. Schoonbeek, G. Yu, E. Baggs, A. I. Witek, I. Yadav, K. V. Krasileva, J. D. G. Jones, C. Uauy, B. Keller, C. J. Ridout, B. B. H. Wulff, The NLR-annotator tool enables annotation of the intracellular immune receptor repertoire. Plant Physiol. 183, 468–482 (2020).

41. M. Maccaferri, N. S. Harris, S. O. Twardziok, R. K. Pasam, H. Gundlach, M. Spannagl, D. Ormanbekova, T. Lux, V. M. Prade, S. G. Milner, A. Himmelbach, M. Mascher, P. Bagnaresi, P. Faccioli, P. Cozzi, M. Lauria, B. Lazzari, A. Stella, A. Manconi, M. Gnocchi, M. Moscatelli, R. Avni, J. Deek, S. Biyiklioglu, E. Frascaroli, S. Corneti, S. Salvi, G. Sonnante, F. Desiderio, C. Marè, C. Crosatti, E. Mica, H. Özkan, B. Kilian, P. De Vita, D. Marone, R. Joukhadar, E. Mazzucotelli, D. Nigro, A. Gadaleta, S. Chao, J. D. Faris, A. T. O. Melo, M. Pumphrey, N. Pecchioni, L. Milanesi, K. Wiebe, J. Ens, R. P. MacLachlan, J. M. Clarke, A. G. Sharpe, C. S. Koh, K. Y. H. Liang, G. J. Taylor, R. Knox, H. Budak, A. M. Mastrangelo, S. S. Xu, N. Stein, I. Hale, A. Distelfeld, M. J. Hayden, R. Tuberosa, S. Walkowiak, K. F. X. Mayer, A. Ceriotti, C. J. Pozniak, L. Cattivelli, Durum wheat genome highlights past domestication signatures and future improvement targets. Nat. Genet. 51, 885–895 (2019).

42. E. D. Badaeva, B. Friebe, B. S. Gill, Genome differentiation in Aegilops. 1. Distribution of highly repetitive DNA sequences on chromosomes of diploid species. Genome. 39, 293–306 (1996).

43. E. D. Badaeva, B. Friebe, B. S. Gill, Genome differentiation in Aegilops. 2. Physical mapping of 5S and 18S-26S ribosomal RNA gene families in diploid species. Genome. 39, 1150–1158 (1996).

44. B. Maestra, T. Naranjo, Homoeologous relationships of Aegilops speltoides chromosomes to bread wheat. Theor. Appl. Genet. 97, 181–186 (1998).

45. H. Ankori, D. Zohary, Natural Hybridization between Aegilops sharonensis and Ae. longissima: A Morphological and Cytological Study. Cytologia (Tokyo). 27, 314–324 (1962).

46. H. Sela, S. Ezrati, P. D. Olivera, Genetic diversity of three Israeli wild relatives of wheat from the Sitopsis section of Aegilops. Isr. J. Plant Sci. 65, 161–174 (2018).

47. E. Millet, Exploitation of Aegilops species of section Sitopsis for wheat improvement. Isr. J. Plant Sci. 55, 277–287 (2007).

48. E. Millet, B. J. Steffenson, R. Prins, H. Sela, A. M. Przewieslik-Allen, Z. A. Pretorius, Genome Targeted Introgression of Resistance to African Stem Rust from Aegilops sharonensis into Bread Wheat. Plant Genome. 10 (2017).

49. E. Millet, J. Manisterskia Distelfeld, J. Deek, a Wan, X. Chen, B.J. Steffenson, Introgression of leaf rust and stripe rust resistance from. Genome. 316, 309–316 (2014).

50. G. Yu, N. Champouret, B. Steuernagel, P. D. Olivera, J. Simmons, C. Williams, R. Johnson, M. J. Moscou, I. Hernández-Pinzón, P. Green, H. Sela, E. Millet, J. D. G. Jones, E. R. Ward, B. J. Steffenson, B. B. H. Wulff, Discovery and characterization of two new stem rust resistance genes in Aegilops sharonensis. Theor. Appl. Genet. 130, 1207–1222 (2017).

51. L. Qiu, N. Liu, H. Wang, X. Shi, F. Li, Q. Zhang, W. Wang, W. Guo, Z. Hu, H. Li, J. Ma, Q. Sun, C. Xie, Fine mapping of a powdery mildew resistance gene MlIW39 derived from wild emmer wheat (Triticum turgidum ssp. dicoccoides). Theor. Appl. Genet. (2021).

52. K. Gaurav, S. Arora, P. Silva, J. Sánchez-Martín, R. Horsnell, L. Gao, G. S. Brar, V. Widrig, J. Raupp, N. Singh, S. Wu, S. M. Kale, C. Chinoy, P. Nicholson, J. Quiroz-Chávez, J. Simmonds, S. Hayta, M. A. Smedley, W. Harwood, S. Pearce, D. Gilbert, N. Kangara, C. Gardener, M. Forner-Martínez, J. Liu, G. Yu, S. Boden, A. Pascucci, S. Ghosh, A. N. Hafeez, J. Waites, J. Cheema, B. Steuernagel, M. Patpour, A. Fejer Justesen, S. Liu, J. C. Rudd, R. Avni, A. Sharon, B. Steiner, R. Pasthika Kirana, H. Buerstmayr, A. A. Mehrabi, F. Y. Nasyrova, N. Chayut, O. Matny, B. J. Steffenson, N. Sandhu, P. Chhuneja, E. Lagudah, A. F. Elkot, S. Tyrrell, X. Bian, R. P. Davey, M. Simonsen, L. Schauser, V. K. Tiwari, H. Randy Kutcher, P. Hucl, A. Li, D.-C. Liu, L. Mao, S. Xu, G. Brown-Guedira, J. Faris, J. Dvorak, M.-C. Luo, K. Krasileva, T. Lux, S. Artmeier, K. F. X Mayer, C. Uauy, M. Mascher, A. R. Bentley, B. Keller, J. Poland, B. B. H Wulff, bioRxiv, in press (available at https://doi.org/10.1101/2021.01.31.428788).

53. A. N. Hafeez, S. Arora, S. Ghosh, D. Gilbert, R. L. Bowden, B. B. H. Wulff, Creation and judicious application of a wheat resistance gene atlas. Mol. Plant (2021).

54. H. B. Zhang, X. Zhao, X. Ding, A. H. Paterson, R. A. Wing, Preparation of megabase-size DNA from plant nuclei. Plant J. 7, 175–184 (1995).

55. S. Walkowiak, L. Gao, C. Monat, G. Haberer, M. T. Kassa, J. Brinton, R. H. Ramirez-Gonzalez, M. C. Kolodziej, E. Delorean, D. Thambugala, V. Klymiuk, B. Byrns, H. Gundlach, V. Bandi, J. N. Siri, K. Nilsen, C. Aquino, A. Himmelbach, D. Copetti, T. Ban, L. Venturini, M. Bevan, B. Clavijo, D. H. Koo, J. Ens, K. Wiebe, A. N’Diaye, A. K. Fritz, C. Gutwin, A. Fiebig, C. Fosker, B. X. Fu, G. G. Accinelli, K. A. Gardner, N. Fradgley, J. Gutierrez-Gonzalez, G. Halstead-Nussloch, M. Hatakeyama, C. S. Koh, J. Deek, A. C. Costamagna, P. Fobert, D. Heavens, H. Kanamori, K. Kawaura, F. Kobayashi, K. Krasileva, T. Kuo, N. McKenzie, K. Murata, Y. Nabeka, T. Paape, S. Padmarasu, L. Percival-Alwyn, S. Kagale, U. Scholz, J. Sese, P. Juliana, R. Singh, R. Shimizu-Inatsugi, D. Swarbreck, J. Cockram, H. Budak, T. Tameshige, T. Tanaka, H. Tsuji, J. Wright, J. Wu, B. Steuernagel, I. Small, S. Cloutier, G. Keeble-Gagnère, G. Muehlbauer, J. Tibbets, S. Nasuda, J. Melonek, P. J. Hucl, A. G. Sharpe, M. Clark, E. Legg, A. Bharti, P. Langridge, A. Hall, C. Uauy, M. Mascher, S. G. Krattinger, H. Handa, K. K. Shimizu, A. Distelfeld, K. Chalmers, B. Keller, K. F. X. Mayer, J. Poland, N. Stein, C. A. McCartney, M. Spannagl, T. Wicker, C. J. Pozniak, Multiple wheat genomes reveal global variation in modern breeding. Nature. 588, 277–283 (2020).

56. S. Beier, A. Himmelbach, C. Colmsee, X. Q. Zhang, R. A. Barrero, Q. Zhang, L. Li, M. Bayer, D. Bolser, S. Taudien, M. Groth, M. Felder, A. Hastie, H. Šimková, H. Staňková, J. Vrána, S. Chan, M. Muñoz-Amatriaín, R. Ounit, S. Wanamaker, T. Schmutzer, L. Aliyeva-Schnorr, S. Grasso, J. Tanskanen, D. Sampath, D. Heavens, S. Cao, B. Chapman, F. Dai, Y. Han, H. Li, X. Li, C. Lin, J. K. McCooke, C. Tan, S. Wang, S. Yin, G. Zhou, J. A. Poland, M. I. Bellgard, A. Houben, J. Doležel, S. Ayling, S. Lonardi, P. Langridge, G. J. Muehlbauer, P. Kersey, M. D. Clark, M. Caccamo, A. H. Schulman, M. Platzer, T. J. Close, M. Hansson, G. Zhang, I. Braumann, C. Li, R. Waugh, U. Scholz, N. Stein, M. Mascher, Construction of a map-based reference genome sequence for barley, Hordeum vulgare L. Sci. Data. 4 (2017).

57. D. Kim, B. Langmead, S. L. Salzberg, HISAT: A fast spliced aligner with low memory requirements. Nat. Methods. 12, 357–360 (2015).

58. M. Pertea, G. M. Pertea, C. M. Antonescu, T. C. Chang, J. T. Mendell, S. L. Salzberg, StringTie enables improved reconstruction of a transcriptome from RNA-seq reads. Nat. Biotechnol. 33, 290–295 (2015).

59. G. Gremme, V. Brendel, M. E. Sparks, S. Kurtz, Engineering a software tool for gene structure prediction in higher organisms. Inf. Softw. Technol. 47, 965–978 (2005).

60. S. Ghosh, C. K. K. Chan, Analysis of RNA-seq data using TopHat and cufflinks. Methods Mol. Biol. 1374, 339–361 (2016).

61. M. Stanke, O. Schöffmann, B. Morgenstern, S. Waack, Gene prediction in eukaryotes with a generalized hidden Markov model that uses hints from external sources. BMC Bioinformatics. 7 (2006).

62. K. J. Hoff, M. Stanke, Predicting Genes in Single Genomes with AUGUSTUS. Curr. Protoc. Bioinforma. 65 (2019), doi:10.1002/cpbi.57.

63. B. J. Haas, S. L. Salzberg, W. Zhu, M. Pertea, J. E. Allen, J. Orvis, O. White, C. R. Robin, J. R. Wortman, Automated eukaryotic gene structure annotation using EVidenceModeler and the Program to Assemble Spliced Alignments. Genome Biol. 9 (2008).

64. B. J. Haas, A. L. Delcher, S. M. Mount, J. R. Wortman, R. K. Smith, L. I. Hannick, R. Maiti, C. M. Ronning, D. B. Rusch, C. D. Town, S. L. Salzberg, O. White, Improving the Arabidopsis genome annotation using maximal transcript alignment assemblies. Nucleic Acids Res. 31, 5654–5666 (2003).

65. M. G. Grabherr, J. Z. Levin, D. A. Thompson, I. Amit, X. Adiconis, L. Fan, R. Raychowdhury, Q. Zeng, Z. Chen, E. Mauceli, N. Hacohen, A. Gnirke, N. Rhind, F. di Palma, B. W. Birren, C. Nusbaum, K. Lindblad-Toh, Full-length transcriptome assembly from RNA-Seq data without a reference genome. Nat. Biotechnol. 29, 644–652 (2011).

66. S. F. Altschul, W. Gish, W. Miller, E. W. Myers, D. J. Lipman, Basic local alignment search tool. J. Mol. Biol. 215, 403–410 (1990).

67. H. Li, Minimap2: Pairwise alignment for nucleotide sequences. Bioinformatics. 34, 3094– 3100 (2018).

68. A. R. Quinlan, I. M. Hall, BEDTools: A flexible suite of utilities for comparing genomic features. Bioinformatics. 26, 841–842 (2010).

69. M. N. Price, P. S. Dehal, A. P. Arkin, FastTree 2 -Approximately maximum-likelihood trees for large alignments. PLoS One. 5 (2010), doi:10.1371/journal.pone.0009490.

70. F. Limin, N. Beifang, Z. Zhengwei, W. Sitao, L. Weizhong, CD-HIT: accelerated for clustering the next generation sequencing data. Bioinformatics. 28, 3150–3152 (2012).

