## Supplementary figures and images for "Genome sequences of *Aegilops* species of section Sitopsis reveal phylogenetic relationships and provide resources for wheat improvement"

### Figure S4

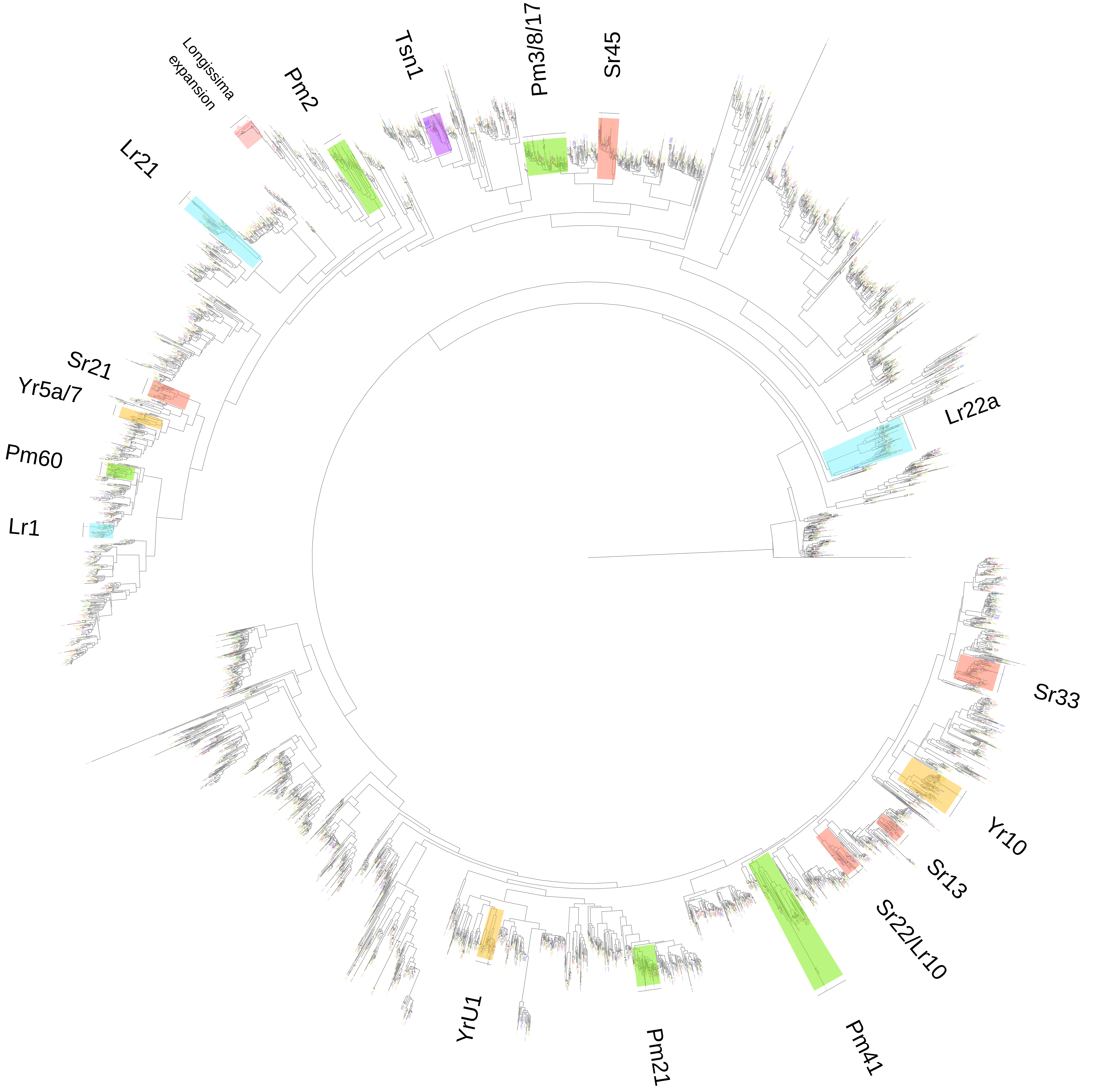
